# Flavonoids as anxiolytics in animal tests: Bibliometry and meta-analysis of the effects of flavonoids on anxiety-like behavior in animal tests

**DOI:** 10.1101/2024.05.27.596117

**Authors:** Jhenify Albuquerque Machado, Danilo Brandão Araújo, Monica Lima-Maximino, Diógenes Henrique de Siqueira-Silva, Bernardo Tomchinsky, Jonathan Cueto-Escobedo, Juan Francisco Rodríguez-Landa, Caio Maximino

## Abstract

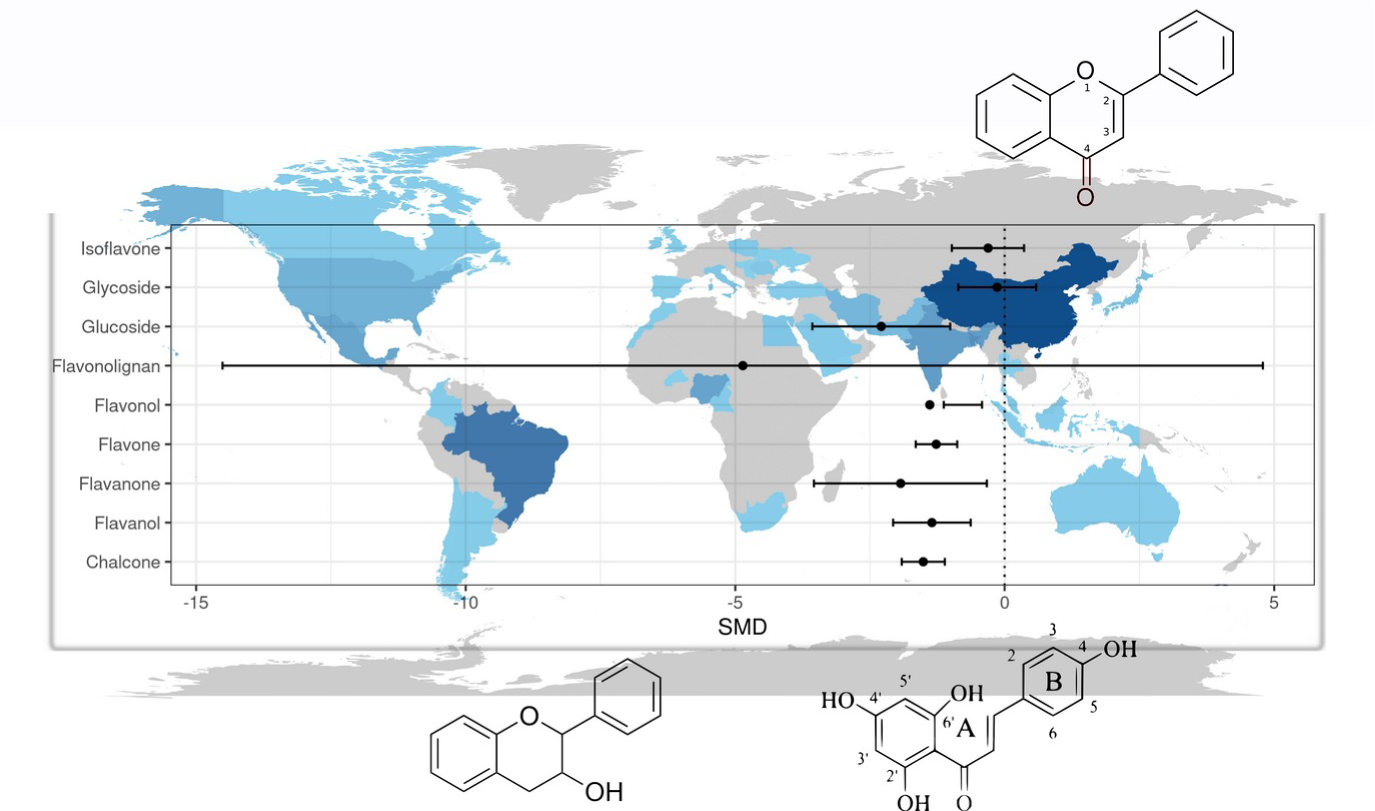

Flavonoids are natural secondary compounds of plants with a basic composition derived from polyphenols that can produce a plethora of different neurochemical effects, some of which are relevant to anxiety disorders. As such, many flavonoids have been evaluated in behavioral screens in preclinical research on anxiolytics. Given the many different molecular subclasses of flavonoids, the many different molecular targets that have been proposed for these compounds, and the different research priorities that arise in preclinical research, we sought to map the potential of flavonoids as anxiolytics by bibliometric analysis and a meta-analysis of animal tests using these compounds. Bibliometric analysis suggest that the field is highly concentrated on few research groups that are mostly located in the Global South, suggesting the need to improve international collaborations. The themes which emerged in the bibliometric analysis are driven by the exploratory steps of pharmacological research, including finding anxioselective effects and looking for dose-response patterns; this suggests that the field, as a whole, could benefit from more mechanistic and confirmatory research. The meta-analysis showed strong evidence for an anxiolytic-like effect of flavonoids on animal tests, including assays made in rats, mice, and zebrafish (SMD = -1.4398, 95%CI[-1.7319, - 1.1477]). Subgroup analysis suggested that this effect is present in acute treatment (SMD = -1.14, 95%CI[-1.36, -0.93]), but not after chronic treatment (SMD = -1.96, 95%CI[-4.93; 1.01]). For all molecule subclasses included in the study, only isoflavones, glycoside derivatives, and flavanolignans did not show evidence of an anxiolytic-like effect, although the low number of studies including these subclasses is low. We finish with a set of recommendations for preclinical research on the anxiolytic potential of flavonoids.

## 1. Introduction

Anxiety disorders such as generalized anxiety disorder, social anxiety disorder, and panic disorder are pathological states characterized by psychological and physical symptoms that severely deteriorates the quality of life of a great percentage of the population around the world (Yang et al., 2021). The first-line treatments against anxiety include psychotherapy and pharmacotherapy based on drugs such as Selective serotonin reuptake inhibitors (SSRIs) and serotonin-norepinephrine reuptake inhibitors (SNRIs), however, benzodiazepines are still used in several countries despite their adverse effects such as pharmacological tolerance, withdrawal symptoms, dependence, sedation, and cognitive impairment (Cueto-Escobedo, García-García, et al., 2020; Gomez et al., 2018).

The need of new treatments without undesired side effects has led to the research of secondary metabolites present in plants, which are first tested at the preclinical level using validated behavioral models of anxiety in animals, such as the elevated plus maze (EPM), the light-dark test (LDT) and the open field test (OFT) among others used with laboratory rodents (Gencturk & Unal, 2024), and the novel tank test (NTT), the light-dark test (LDT) and others used in the zebrafish (Gerlai, 2023; Kysil et al., 2017). These tests show great utility as behavioral screens to explore the anxiolytic-like effects of plant secondary metabolites, such as flavonoids (Cueto-Escobedo et al., 2022; German-Ponciano et al., 2018).

Flavonoids are natural secondary compounds of plants with a basic composition derived from polyphenols (hydroxyls attached to phenolic rings) with low molecular weight. Widely found in higher plants, there are around six thousand flavonoid molecules described in nature, which can be categorized into groups according to their chemical structures, size, and the degree of oxidation of the phenol, including chalcones, flavones, flavanones, flavonols, dihydroflavonols, isoflavones, anthocyanins, and aurones (Panche et al., 2016). In plants, they are synthesized through the shikimic acid or acetate (acetyl coenzyme A) metabolic pathways, and the formation of a chalcone that is the precursor of all flavonoids. For plants, flavonoids have adaptive importance as antioxidants, in hormone production, communication, allelopathy, attracting pollinators and dispersers, protection against herbivory, antimicrobial activity, or protection against freezing.

The presence of flavonoids in plants is related to genetic, environmental, and cultural (cultivation) factors, such as chemotypes, circadian cycles, phenology and maturation, part and structure of the plant, solar radiation, soil characteristics, water availability, altitude, temperature, among others. Another relevant factor in the production and availability of flavonoids, considering metabolic pathways, plant morphology and anatomy, environmental issues, and ecological functions, is the storage of these substances. They can be concentrated in fruits, flowers, barks, or in specialized structures such as trichomes, glands, and resin ducts (Samantha et al., 2011).

It is important to note that in each species, more than a dozen types of flavonoids are synthesized, usually with a predominance of a few that are used as molecular markers and generally are those with the greatest pharmacological relevance. Considering the restricted occurrence of each group of secondary compounds in the plant kingdom, flavonoids are relevant molecules for studies in plant chemotaxonomy (Cronquist system), and pharmacology. Some groups of phenols are related to specific families or genera of plants (Alkin, 2017).

Despite being found in low concentrations in plants, flavonoids have significant applications for the human population, such as dyes, food, nutraceuticals, cosmetics, flavorings, and medicines. They are the predominant component in many medicinal plants known (Alkin, 2017). In pharmacological use, following the diversity of flavonoids, they are associated with a wide range of medical treatments such as antioxidant, anti-inflammatory, anticancer, effects on the central nervous system, coronary diseases, among others (Abou Baker, 2022; Panche et al., 2016; Tungmunnithum et al., 2018). Flavonoids are of special interest to psychopharmacology, given that they are major components of medicinal plants with possible anxiolytic effect in different cultures (German-Ponciano et al., 2018; Z. Liu et al., 2021; López-Rubalcava & Estrada-Camarena, 2016; Rodrigues et al., 2008). Experimental results show that flavonoids and their conjugates derivatives may cross the bloodbrain barrier and exert pharmacological action on the central nervous system (Jäger & Saaby, 2011; Youdim et al., 2003) by activating neuronal signaling pathways (Spencer, 2007) and neurotransmitter systems (i.e., serotonergic, GABAergic, noradrenergic, and dopaminergic, among others) involved in neuropsychiatric disorders as anxiety and depression (German-Ponciano et al., 2018; C. Wang et al., 2023; H. Wang et al., 2023).

There is considerable evidence that some naturally occurring flavonoids are ligands at the central benzodiazepine receptor (i.e., the benzodiazepine site at GABA_A_ receptors)(Hanrahan et al., 2011; Marder et al., 2001). A pharmacophore model suggests that flavone derivatives bind the same site of benzodiazepines (BZDs), due mainly to the negatively charged oxygen atom of the carbonyl group of the flavonoids and with the nature of the substituent in position 3’ (Marder et al., 2001). Kahnberg et al. (2002) produced a pharmacophore model with two additional regions of steric repulsive interactions between flavone ligands and the BZD site. Flavones, especifically, appear to produce a range of biological activities at BZD sites, with some molecules acting as full positive allosteric modulators, others as partial positive allosteric modulators, and othes as negative modulators (Hanrahan et al., 2011; Wasowski & Marder, 2012). Interestingly, flavones as apigenin, and hesperidin target on the GABAA/benzodiazepine receptor complex producing anxiolytic-like effects, however, depending on the doses also may produce sedative and sleep-enhancing properties (Marder et al., 2003). Chrysin is another flavone that target on the GABA_A_/benzodiazepine receptor complex producing anxiolytic-like effects, but differently to hesperidin and apigenin this flavone is devoid of sedative and motor effects (Cueto-Escobedo, Andrade-Soto, et al., 2020; Wolfman et al., 1994) – an advantage compared with other flavonoid and benzodiazepines. This also highlights that flavonoids may differ on the neurochemical and neurotrophic pathways contributing to their anxiolytic-like effects.

Flavonoids interact with neurotransmission systems and neurotrophic factors and produce antioxidant and anti-inflammatory effects in the brain, contributing to their anxiolytic-like effects (Rodríguez-Landa, German-Ponciano, et al., 2022). For example, it has been identified that 2.5 and 5 mg/kg of apigenin isolated from *Stachys tibetica* produced anxiolytic-like effects in male and female Wistar rats evaluated in the EPM (Kumar & Bhat, 2014). The effects of apigenin might be associated with action on the GABA_A_/benzodiazepine receptor complex and the 5-HT_1A_/5-HT_2A_ receptors (Amin et al., 2022; Salgueiro et al., 1997). On the other hand, 50 mg/kg/28 days hesperidin reduces anxiety-like behavior in the EPM in mice treated with the dopaminergic neurotoxin 6-OHDA, without significant changes in spontaneous locomotor activity in the OFT, which was related with attenuation of proinflammatory cytokines tumor necrosis factor-α, interferon gamma, interleukin-1β, interleukin-2, and interleukin-6; in addition to increases in the levels of neurotrophic factors, including neurotrophin-3, brain-derived neurotrophic factor and nerve growth factor in the striatum and other brain structures (Antunes et al., 2020). Similarly, 25, 50 and 100 mg/kg chrysin produces anxiolytic-like effects in male Sprague-Dawley rats evaluated in the LDT, in a similar fashion that apigenin, through actions on the GABA_A_ receptor (Zanoli et al., 2000); while 2, 4, and 8 μmol/kg produces anxiolytic-like effects associated with an increases of the number of Fos-immunoreactive cells in the lateral septal nucleus (Germán-Ponciano et al., 2020). It is important to mention that the anxiolytic-like effects of chrysin are depending on the ovarian cycle phase in female Wistar rats. The administration of chrysin produces anxiolytic-like effects during metestrus-diestrus phase and devoid of effects during proestrus-estrus phase, those effects were blocked for the non-selective GABA_A_ receptor antagonist picrotoxin (Rodríguez-Landa et al., 2021). However, when chrysin is microinjected into the dorsal hippocampus anxiolytic-like effects were detected during metestrus-diestrus phase, but anxiogenic-like effects were produced during proestrus-estrus phase, a similar effect to that produced by the neurosteroid allopregnanolone (Rodríguez-Landa, Hernández-López, et al., 2022). This shows that the effects of flavonoids may vary due to pharmacological interactions with steroid hormones and possibly with other substances.

These effects of flavonoids suggest that this class of molecules can produce important effects on anxiety and defensive behavior. Indeed, a recent meta-analysis of small clinical studies suggested that flavonoids produce significant anxiolytic effects (Jia et al., 2023). However, not only do flavonoids considerably differ in terms of their actions on the central BZD site (Hanrahan et al., 2011; Wasowski & Marder, 2012), these molecules also target other receptors, and present receptor-independent mechanisms such as reactive oxygen species (ROS) sequestration. Thus, the field of preclinical research on flavonoids as anxiolytics is wide, not only due to the many different possible mechanisms of action of specific molecule subclasses, but also due to different research priorities within the field. The present work attempted to map this variation through both a bibliometric analysis of the field, trying to discern the conceptual and social structure of the field of preclinical research on flavonoids as anxiolytics, and a meta-analysis of animal experiments on anxiolytic-like effects of these molecules.

## 2. Methods

### 2.1. Bibliometric analysis

#### 2.1.1. Data collection and screening

Data retrieval for bibliometric analysis was conducted on 18^th^ May, 2024, using PubMed (https://www.ncbi.nlm.nih.gov/pubmed) to search for articles describing the effects of flavonoids on anxiety-like behavior in animal tests. The following string was used: ((((((((flavonoid[Title/Abstract]) AND (elevated plus-maze[Title/Abstract])) OR ((flavonoid[Title/Abstract]) AND (light-dark test[Title/Abstract]))) OR ((flavonoid[Title/Abstract]) AND (open-field test[Title/Abstract]))) OR ((flavonoid[Title/Abstract]) AND (holeboard[Title/Abstract]))) OR ((flavonoid[Title/Abstract]) AND (zebrafish[Title/Abstract]) AND (anxiety[Title/Abstract]))) OR (((flavonoid[Title/Abstract]) AND (rat[Title/Abstract]) AND (anxiety[Title/Abstract])))) OR (((flavonoid[Title/Abstract]) AND (mouse[Title/Abstract]) AND (anxiety[Title/Abstract])))) OR (((flavonoid[Title/Abstract]) AND (mice[Title/Abstract]) AND (anxiety[Title/Abstract]))). A search filter optimized to finding studies on animal experimentation on PubMed was used (Hooijmans et al., 2010). Only documents written in English were retained. A total of 197 documents were retained for bibliometric analysis.

#### 2.1.2. Bibliometric and scientometric analyses

Bibliometric analyses were made using the R package ‘bibliometrix’ (Aria & Cuccurullo, 2017). The package was used first to conduct a descriptive analysis, quantifying timespan, number of documents, annual scientific production, annual growth rate, document average age, average citations per document, description of most relevant sources and affiliations, and analyses of corresponding author’s countries. Influential sources were analyzed using Bradford’s law (Bradford, 1934/1985). The conceptual structure of the field was analyzed using co-word analysis through keyword co-occurrences; a network layout was generated using the Fruchterman-Reingold algorithm, plotting the main 50 cited references, with vertex similarities normalized using association strength. This network was described in terms of size, density, transitivity, degree of centralization, and average path length. A thematic map was drawn by crossing the density and centrality of keyword cooccurrences, placing themes in four quadrants: (1) upper-right quadrant: motor-themes (themes that are both well-developed and important for the structure of the specific research field); (2) lower-right quadrant: basic themes (themes that are important for the research field but underveloped); (3) lower-left quadrant: emerging or disappearing themes (themes that are both weakly developed and with low external ties to other themes); (4) upper-left quadrant: very specialized/niche themes (themes that have well-developed internal ties but low external ties to other themes). Finally, the social structure of the field was analyzed using collaboration networks.

### 2.2. Meta-analysis

#### 2.2.1. Search parameters and data extraction

In order to find and filter data for the meta-analysis, we searched PubMed (https://www.ncbi.nlm.nih.gov/pubmed) for articles describing the effects of flavonoids on anxiety-like behavior in animal tests. The following string was used: ((((((((flavonoid[Title/Abstract]) AND (elevated plus-maze[Title/Abstract])) OR ((flavonoid[Title/Abstract]) AND (light-dark test[Title/Abstract]))) OR ((flavonoid[Title/Abstract]) AND (open-field test[Title/Abstract]))) OR ((flavonoid[Title/Abstract]) AND (holeboard[Title/Abstract]))) OR ((flavonoid[Title/Abstract]) AND (zebrafish[Title/Abstract]) AND (anxiety[Title/Abstract]))) OR (((flavonoid[Title/Abstract]) AND (rat[Title/Abstract]) AND (anxiety[Title/Abstract])))) OR (((flavonoid[Title/Abstract]) AND (mouse[Title/Abstract]) AND (anxiety[Title/Abstract])))) OR (((flavonoid[Title/Abstract]) AND (mice[Title/Abstract]) AND (anxiety[Title/Abstract]))). A search filter optimized to finding studies on animal experimentation on PubMed was used (Hooijmans et al., 2010). The following inclusion criteria were used: studies which included primary behavioral data obtained in tests for anxiety-like behavior in rats, mice, or zebrafish (light/dark test, novel tank test, elevated-plus maze, open-field test, or holeboard test); reporting of appropriate controls; reporting of at least sample sizes and summary statistics (mean and standard deviation or standard error of the mean), either in tables or graphs, for control and treated groups (da Silva Chaves et al., 2018). Whenever an experiment evaluated the effects of other drugs (e.g., GABA_A_ receptor antagonists, to assess the role of these receptors in the effects of the flavonoid) or interventions (e.g, acute stress) on anxiety-like behavior, only control and flavonoid groups were considered, and data under intervention effects was not analyzed. Possible confounds in relation to the role of development were reduced by excluding studies which were not performed on adult animals.

The following data were extracted from each included study: identification (DOI, authors, publication year); species; strain/phenotype; behavioral test that was used; molecule; dose of the flavonoid; means and standard deviations; and sample sizes (*N*) for each group. In studies in which data was represented graphically, means and standard deviations were extracted from figures using the PlotDigitizer online app (https://plotdigitizer.com/app). When multiple dependent variables were reported, only the primary endpoint was used. Following Vesterin et al. (2014), when studies contained multiple comparisons with the control group (e.g., studies with multiple doses of the flavonoid of interest), the sample size for the control group was corrected for multiple comparisons by dividing the number reported by the number of treatment groups.

#### 2.2.2. Statistical analysis

A general meta-analysis was made on all data, and subset analyses were made on data from different molecule class and treatment durations; in the subset analyses, the procedures were the same as those used for the general meta-analysis. The analysis was carried out using the standardized mean difference (SMDs) as the outcome measure (*y_i_*). Qualitative interpretations of the magnitudes of SMDs followed Cohen’s (1988) guidelines. A random-effects model was fitted to the data. The amount of heterogeneity (i.e., tau²), was estimated using the restricted maximum-likelihood estimator (Viechtbauer, 2005). In addition to the estimate of tau², the Q-test for heterogeneity (Cochran, 1954) and the I² statistic were reported. In case any amount of heterogeneity is detected (i.e., tau² > 0, regardless of the results of the Q-test), a prediction interval for the true outcomes was also provided. Studentized residuals and Cook’s distances were used to examine whether studies may be outliers and/or influential in the context of the model. Studies with a studentized residual larger than the 100×(1−0.05 /2×*k*) ^th^ percentile of a standard normal distribution were considered potential outliers (i.e., using a Bonferroni correction with two-sided alpha = 0.05 for *k* comparisons included in the meta-analysis). Studies with a Cook’s distance larger than the median plus six times the interquartile range of the Cook’s distances were considered to be influential. To analyze publication bias, the rank correlation test (Begg & Mazumdar, 1994) and the regression test (Egger et al., 1997), using the standard error of the observed outcomes as predictor, were used to check for funnel plot asymmetry. Meta-analyses were made using R (version 4.1.2), with the package ‘metafor’ (version 3.8-1; Viechtbauer, 2010).

### 2.3. Dose-response curves

Dose-response curves for each of the molecule classes were obtained by fitting generalized log-logistic 5 parameter models (LL5), with dose as independent variable and observed effect size *y_i_* as dependent variable – with the exception of data on chalcones and flavanols, since visual inspection of plots suggested an inverted-U shaped curve; in this case, a 5-parameter Brain-Cousens model was fitted (Brain & Cousens, 1989). To compare the general potency of molecule classes, effective doses (ED_50_) were calculated for each curve. Curves were estimated using R (version 4.1.2), with the package ‘drc’ (version 3.0-1; Ritz et al., 2015).

## 3. Results

### 3.1. Bibliometric analysis

A total of 197 documents were retrieved for the analysis. The documents retrieved for bibliometric analysis spanned 30 years, from 1994 to 2024. The annual growth rate of the field was 8.32% (Figure 2A), and the average document age was 6.6 years. Scientific production showed two peaks: between 2008 an 2010, and again following 2018 and 2019. The average number of co-authors per document was 6.75, with an average of 0.182 documents per author. International co-authorships comprised 17.26% of the documents. With the exception of USA, which was the 7^th^ most productive country in the field, the 10 most productive countries in the field were from the Global South (Figure 2B). A network of keyword co-occurrences (Figure 2C) produced 960 nodes, with a density of 0.023, transitivity of 0.259, a low degree of centralization (0.399), and long average path lengths (2.421). The network revealed two main clusters: the green cluster, representing basic methods and mechanisms (serotonin metabolism, hippocampus); and the purple cluster, describing behavioral pharmacological methods. Thematic mapping (Figure 2D) suggested basic themes that describe basic methods (e.g., “behavior, animal/drug effects” or “dose-response relationship, drug”); motor themes include methods related to phytochemistry and pharmacognosia (e.g., “chromatography, high pressure liquid” and “plant extracts/chemistry/pharmacology”); niche themes include molecular docking, signal transduction mechanisms, and neuropathology; and emerging themes include catechins and inflammation. An analysis of author collaboration networks suggested little collaboration outside of a given research group: the network was sparse (density of 0.007) and non-centralized (centralization of 0.031), with low transitivity (0.787).

**Figure 1.** PRISMA flow diagram of the document screening process for the meta-analysis.

**Figure 2.**
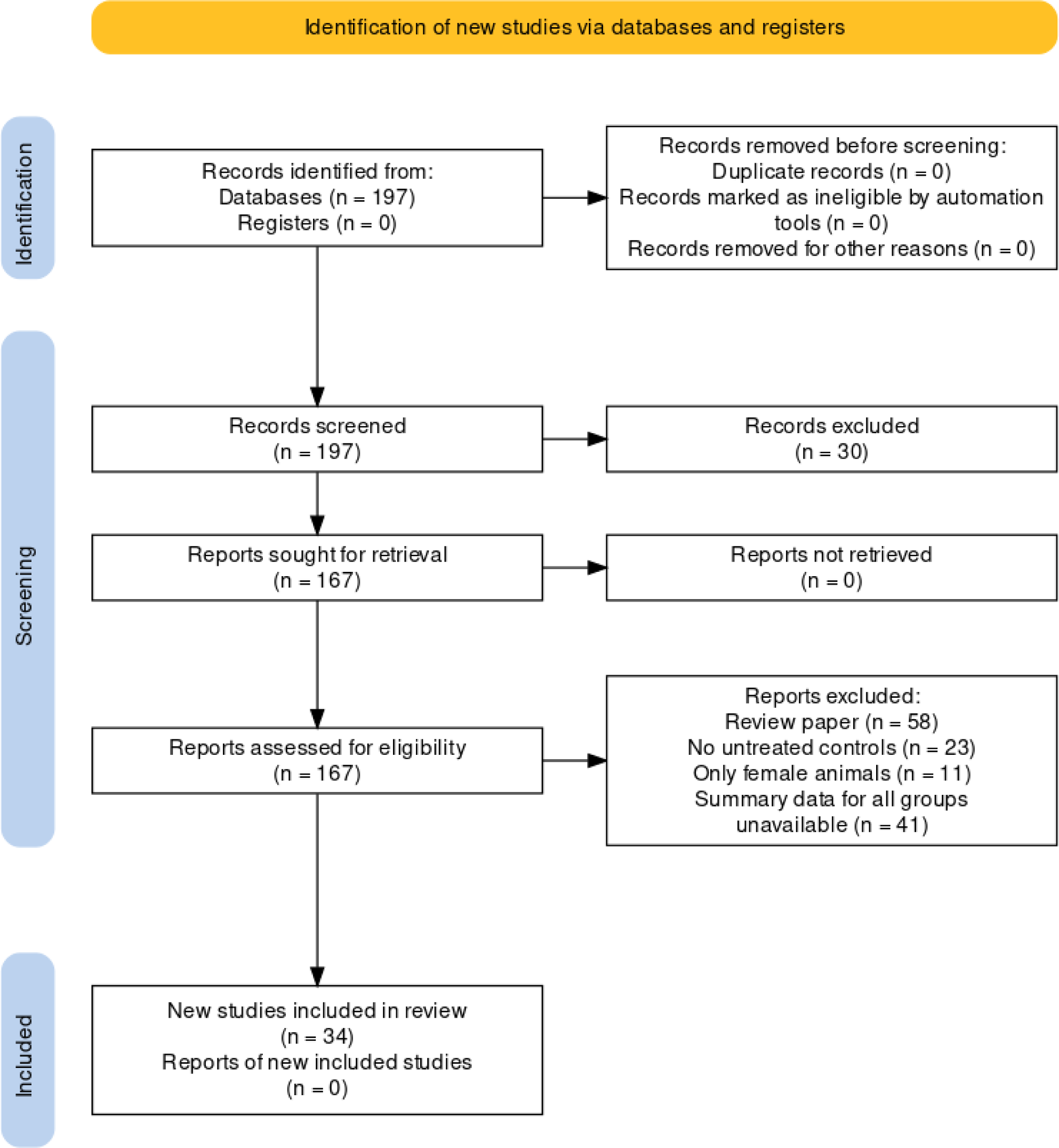
Bibliometric analysis of the field of preclinical research on the anxiolytic-like effects of flavonoids in animal tests. (A) The changes in annual publications from 1994 to 2024. (B) Country scientific production and collaboration status; number of publications per country, either as single-country publications (blue, SCP) or multiple country publications (red, MCP). (C) Keyword co-occurrence network and clusters identified with bibliometrix. Each node represents a keyword that meet the filtering thresholds; node size is correlated with their publication numbers and the curved line represents the co-occurrence relationship between keywords. (D) Thematic map, by density and centrality, based on author keywords.

### 3.2. Meta-analysis

The search parameters retrieved 196 articles in the PubMed database, of which 34 were retained for analysis after application of inclusion criteria. A total of *k* = 172 comparisons were included in the analysis; this is due to the fact that most articles included more than one dose and/or more than one flavonoid. The smallest number of comparisons in a single article was 1, and the largest was 16. A total of 40 different molecules were included in the analysis (Table 1).

**Table 1.**
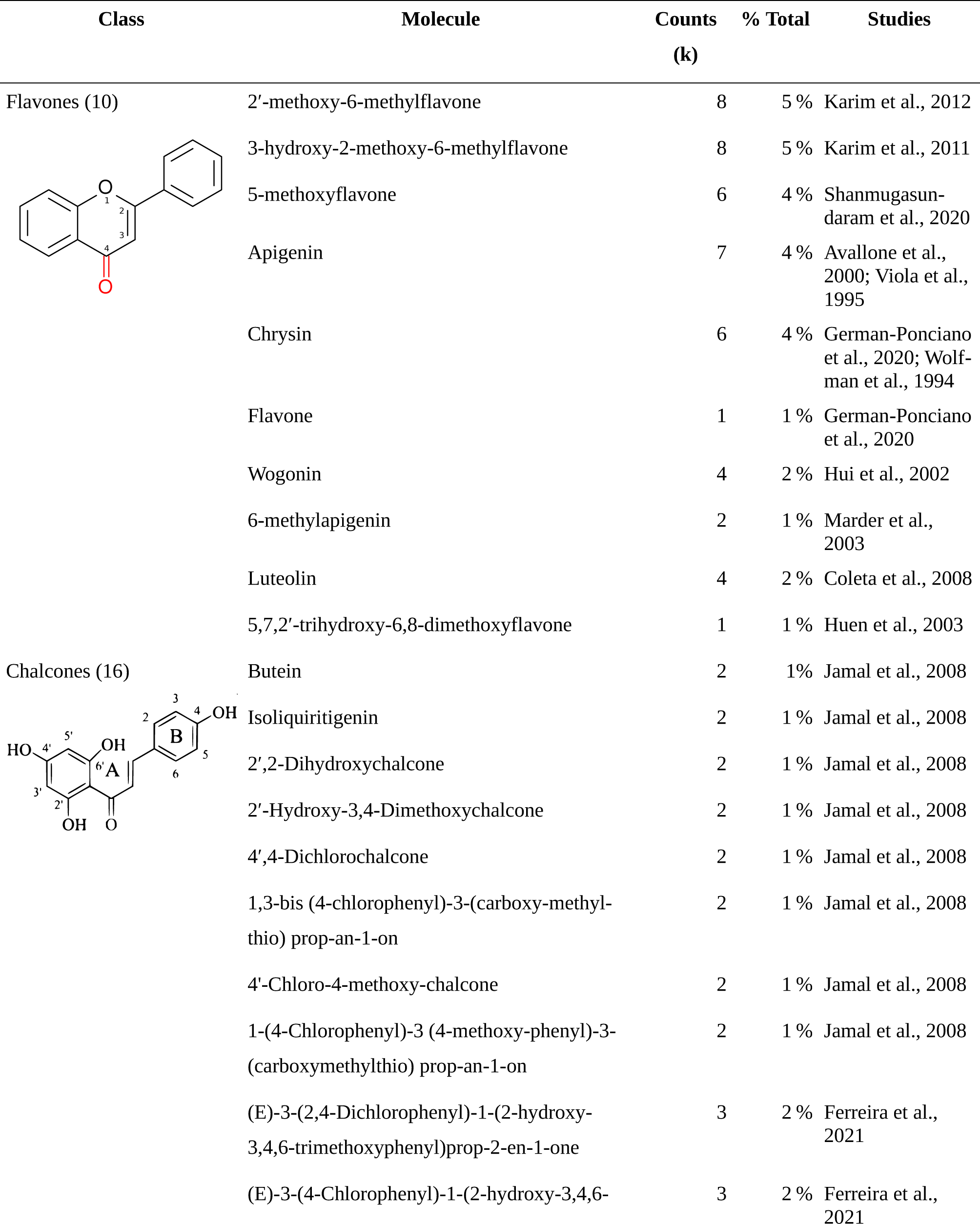
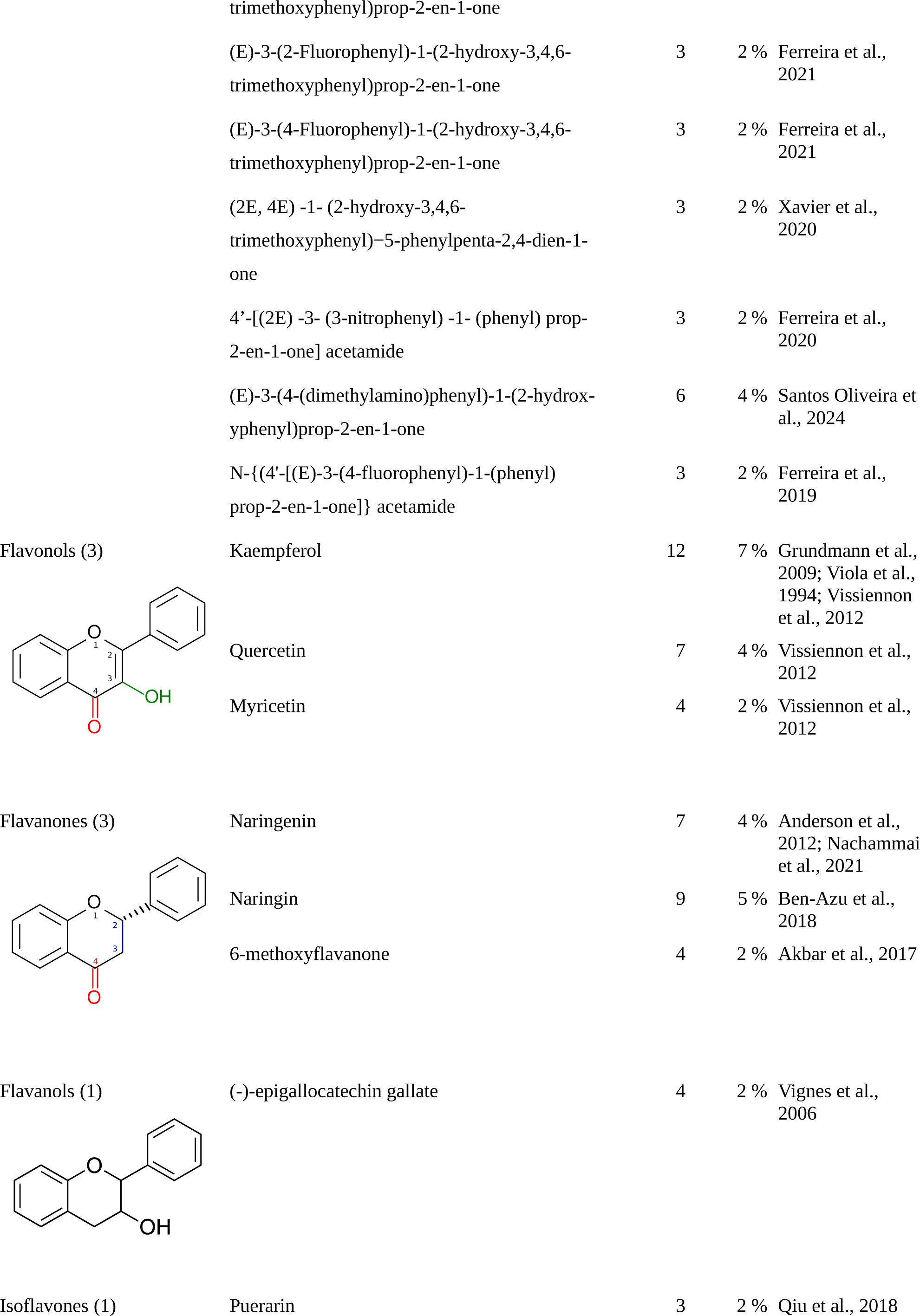
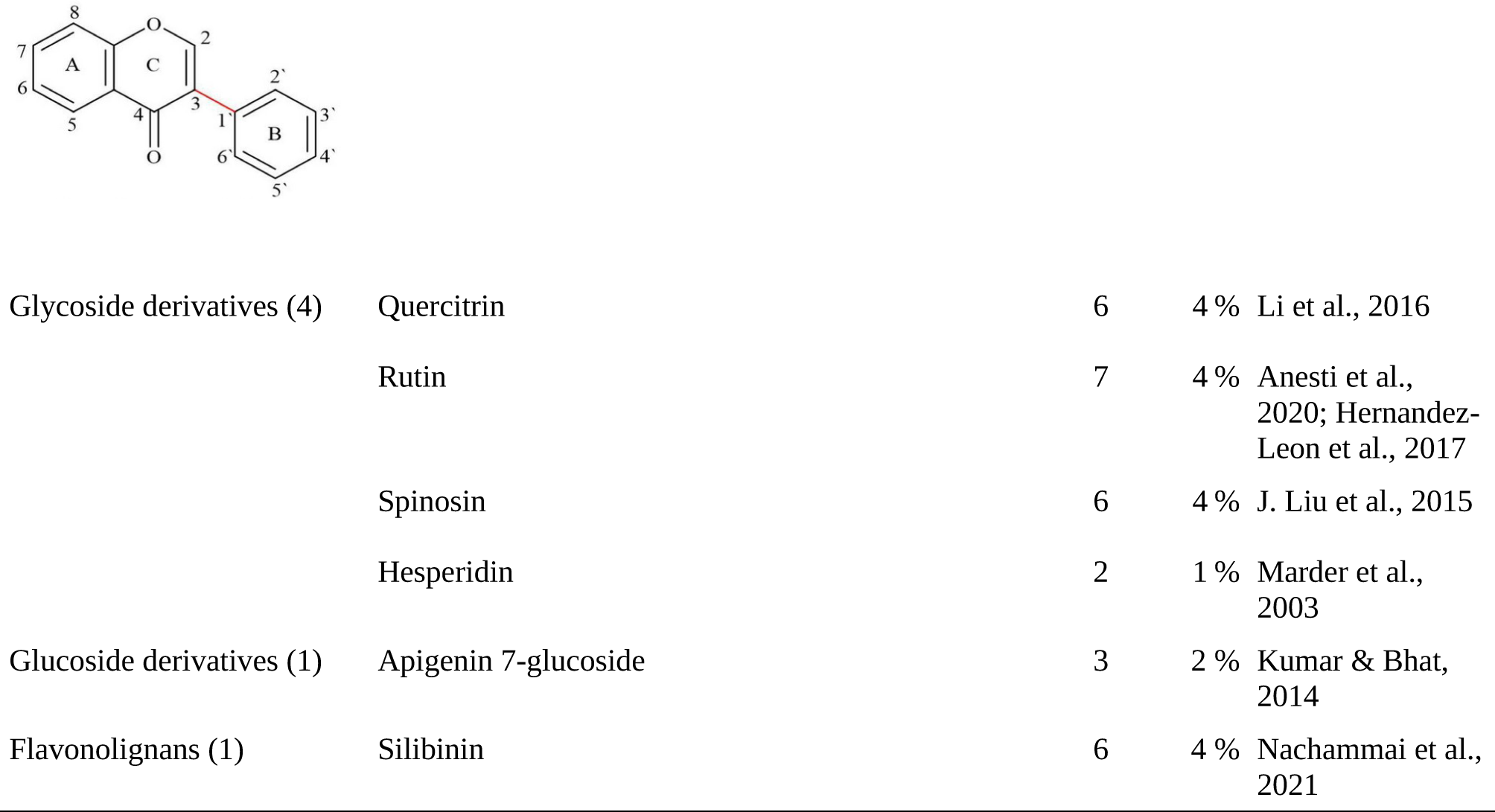
Summary of the studies included in the meta-analysis, including molecule class, specific molecules under the class, total number of comparisons included in the analysis (*k*), proportion of the total comparisons comprised by that molecule, and the reference. A single study can count a large number of comparisons due to the use of a larger number of doses.

For the overall meta-analysis, the observed standardized mean differences ranged from - 20.1471 to 12.9258, with the majority of estimates (87%) being negative in relation to control groups (i.e., suggesting decreased anxiety-like behavior). The estimated average standardized mean difference based on the random-effects model was 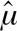 = -1.4398 (95% CI: -1.7319 to -1.1477) (Figure S1), interpreted as a large effect size. Therefore, the average outcome differed significantly from zero (z = -9.6613, p < 0.0001). According to the Q-test, the true outcomes appear to be heterogeneous (Q_[df= 169]_ = 728.8937, p < 0.0001, τ² = 2.7988, I² = 83.1004%). A 95% prediction interval for the true outcomes is given by -4.7318 to 1.8521. Hence, although the average outcome is estimated to favor the flavonoid, in some studies the true outcome may in fact favor the control. An examination of the studentized residuals revealed that several comparisons (the effect of the highest dose of hesperidin on the holeboard in mice in Marder et al., 2003; the effect of the highest dose of silibinin and naringenin on the novel tank test and of the lowest doses of both molecules on the light/dark test in zebrafish in Nachammai et al., 2021) had values larger than ± 3.6204 and may be potential outliers in the context of this model. According to the Cook’s distances, several comparisons (the effects of all doses of silibinin on the novel tank test and of the highest dose of this molecule on the light/dark test in zebrafish in Nachammai et al., 2021) could be considered to be overly influential. The rank correlation and the regression tests indicated potential funnel plot asymmetry (p < 0.001 for all tests; Figure 3), suggesting publication bias towards positive results.

**Figure 3.** Funnel plot for the general meta-analysis. Each plotted point represents the standardized mean difference (SMD) and the square root of the sample size between controls and flavonoid-treated animals for a single comparison. The vertical line represents the average SMD of -1.4398 found in the meta-analysis. The asymmetry of the data points towards the left of this line is suggestive of publication bias.

#### 3.2.1. Subgroup analysis: Treatment duration

Both acute (*k* = 150) and chronic (*k* = 22) treatment regimes were found in the literature search. Acute treatments (Figure S2) appeared to produce consistent anxiolytic-like effects; the estimated average standardized mean difference based on the random-effects model was 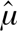 = -1.14 (95% CI: -1.36 to -0.93), suggesting large effects. The average outcome significantly differed from zero (z = -10.305, p < 0.0001). According to the Q-test, the true outcomes appear to be moderately heterogeneous (Q_[df= 149]_ = 400.943, p < 0.0001, τ² = 0.9599, I² = 61.82%).

Chronic treatment, on the other hand, did not produce either anxiolytic-like nor anxiogenic-like effects (Figure S3). The estimated average standardized mean difference based on the random-effects model was 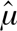 = -1.96 (95% CI: -4.93 to 1.01). The average outcome did not significantly differ from zero (z = -1.29, p =0.195). According to the Q-test, the true outcomes appear to be highly heterogeneous (Q_[df= 21]_ = 162.96, p < 0.0001, τ² = 46.21, I² = 98.55%).

#### 3.2.2. Subgroup analyses: Molecule class

For flavones (*k* = 49), the estimated average standardized mean difference based on the random-effects model was 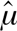 = -1.27 (95% CI: -1.65 to -0.88)(Figure 4A), interpreted as a large effect size. Therefore, the average outcome differed significantly from zero (z = -6.4534, p < 0.0001), suggesting an overall anxiolytic-like effect. According to the Q-test, the true outcomes appear to be heterogeneous (Q_[df= 48]_ = 164.407, p < 0.0001, τ² = 1.0735, I² = 69.84%). There is some evidence for a dose-dependent effect, with higher doses producing larger effects (Figure 4B), with a calculated ED_50_ of 9.0942 ± 3.02 mg/kg (mean ± SEM).

**Figure 4.**
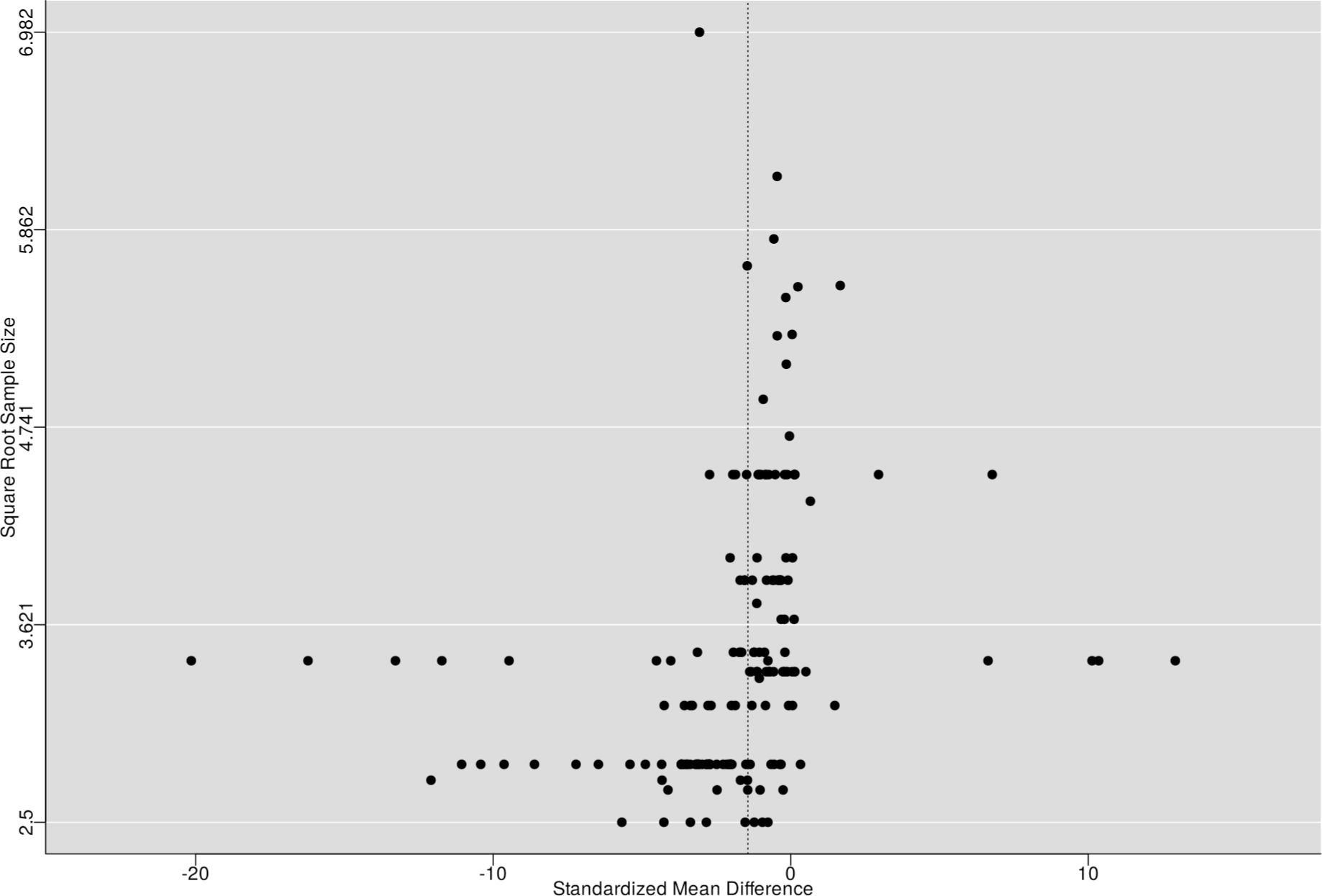
Subgroup analysis for flavones. (A) Forest plot showing the results of 49 comparisons examining the effect of a flavone on anxiety-like behavior in animl tests. The figure shows the standardized mean difference (SMD) between control and flavone-treated groups with corresponding 95% confidence intervals in the individual comparison, based on a random-effects model. A negative standardized mean difference (SMD) corresponds to decreased anxiety-like behavior, while a positive SMD corresponds to increased anxiety-like behavior after flavone treatment. The overall effect size is denoted by the diamond symbol. (B) Estimated dose-response curve, based on the observed effect size (SMD) with increasing flavone dose. Each point corresponds to a single comparison from the forest plot. The curve was obtained by fitting generalized log-logistic 5 parameter models, with dose as independent variable and observed effect size SMD as dependent variable.

Similar results were found for chalcones (*k* = 43), in which the estimated average standardized mean difference based on the random-effects model was 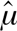 = -1.51 (95% CI: -1.91 to -1.11) (Figure 5A), interpreted as a large effect size. Therefore, the average outcome differed significantly from zero (z = -7.4591, p < 0.0001), suggesting an overall anxiolytic-like effect. According to the Q-test, the true outcomes appear to be heterogeneous (Q_[df= 42]_ = 69.1009, p = 0.005, τ² = 0.478, I² = 28.15%). There is some evidence for a dose-dependent effect, with smaller and larger doses producing larger effects (Figure 5B), with a calculated ED_50_ of 27.127 ± 5.74 mg/kg (mean ± SEM).

**Figure 5.** Subgroup analysis for chalcone. (A) Forest plot showing the results of 43 comparisons examining the effect of a chalcone on anxiety-like behavior in animal tests. The figure shows the standardized mean difference (SMD) between control and chalcone-treated groups with corresponding 95% confidence intervals in the individual comparison, based on a random-effects model. A negative standardized mean difference (SMD) corresponds to decreased anxiety-like behavior, while a positive SMD corresponds to increased anxiety-like behavior after chalcone treatment. The overall effect size is denoted by the diamond symbol. (B) Estimated dose-response curve, based on the observed effect size (SMD) with increasing chalcone dose. Each point corresponds to a single comparison from the forest plot. The curve was obtained by fitting a 5-parameter Brain-Cousens model, with dose as independent variable and observed effect size SMD as dependent variable.

Flavonols (*k* = 23) also appeared to produce an anxiolytic-like effect (Figure 6A). The estimated average standardized mean difference based on the random-effects model was 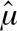 = -1.39 (95% CI: -1.13 to -0.42), interpreted as a medium-to-large effect size. Therefore, the average outcome differed significantly from zero (z = -4.3281, p < 0.0001), suggesting an overall anxiolytic-like effect. According to the Q-test, the true outcomes appear to be heterogeneous (Q_[df= 22]_ = 36.3998, p = 0.0275, τ² = 0.2743, I² = 38.9%). There is some evidence for a dose-dependent effect, with higher doses producing larger effects (Figure 6B), and a calculated ED_50_ of 17.563 ± 7.08 mg/ kg (mean ± SEM).

**Figure 6.**
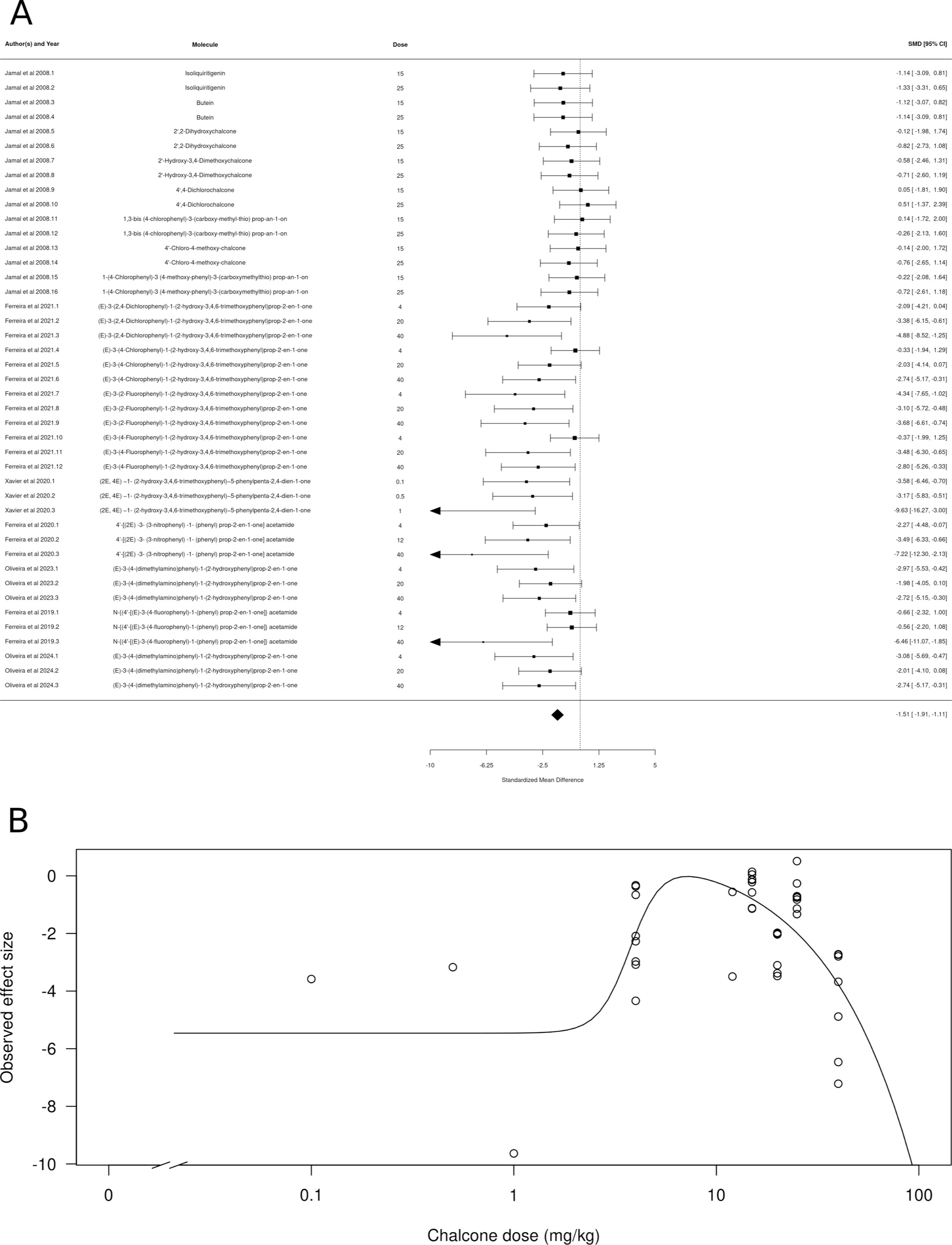
Subgroup analysis for flavonols. (A) Forest plot showing the results of 23 comparisons examining the effect of a flavonol on anxiety-like behavior in animal tests. The figure shows the standardized mean difference (SMD) between control and flavonol-treated groups with corresponding 95% confidence intervals in the individual comparison, based on a random-effects model. A negative standardized mean difference (SMD) corresponds to decreased anxiety-like behavior, while a positive SMD corresponds to increased anxiety-like behavior after flavonol treatment. The overall effect size is denoted by the diamond symbol. (B) Estimated dose-response curve, based on the observed effect size (SMD) with increasing flavonol dose. Each point corresponds to a single comparison from the forest plot. The curve was obtained by fitting generalized log-logistic 5 parameter models, with dose as independent variable and observed effect size SMD as dependent variable.

Flavanones (*k* = 20) also appeared to produce an anxiolytic-like effect (Figure 7A). The estimated average standardized mean difference based on the random-effects model was 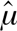 = -1.93 (95% CI: -3.54 to -0.33), interpreted as a small-to-large effect size. Therefore, the average outcome differed significantly from zero (z = -2.363, p = 0.018), suggesting an overall anxiolytic-like effect. According to the Q-test, the true outcomes appear to be highly heterogeneous (Q _[df= 19]_ = 91.4232, p < 0.0001, τ² = 0.2743, I² = 89.61%). There is some evidence for a dose-dependent effect, with higher doses producing slightly larger effects (Figure 7B), and a calculated ED_50_ of 1.02 ± 1.01 mg/kg (mean ± SEM).

**Figure 7.**
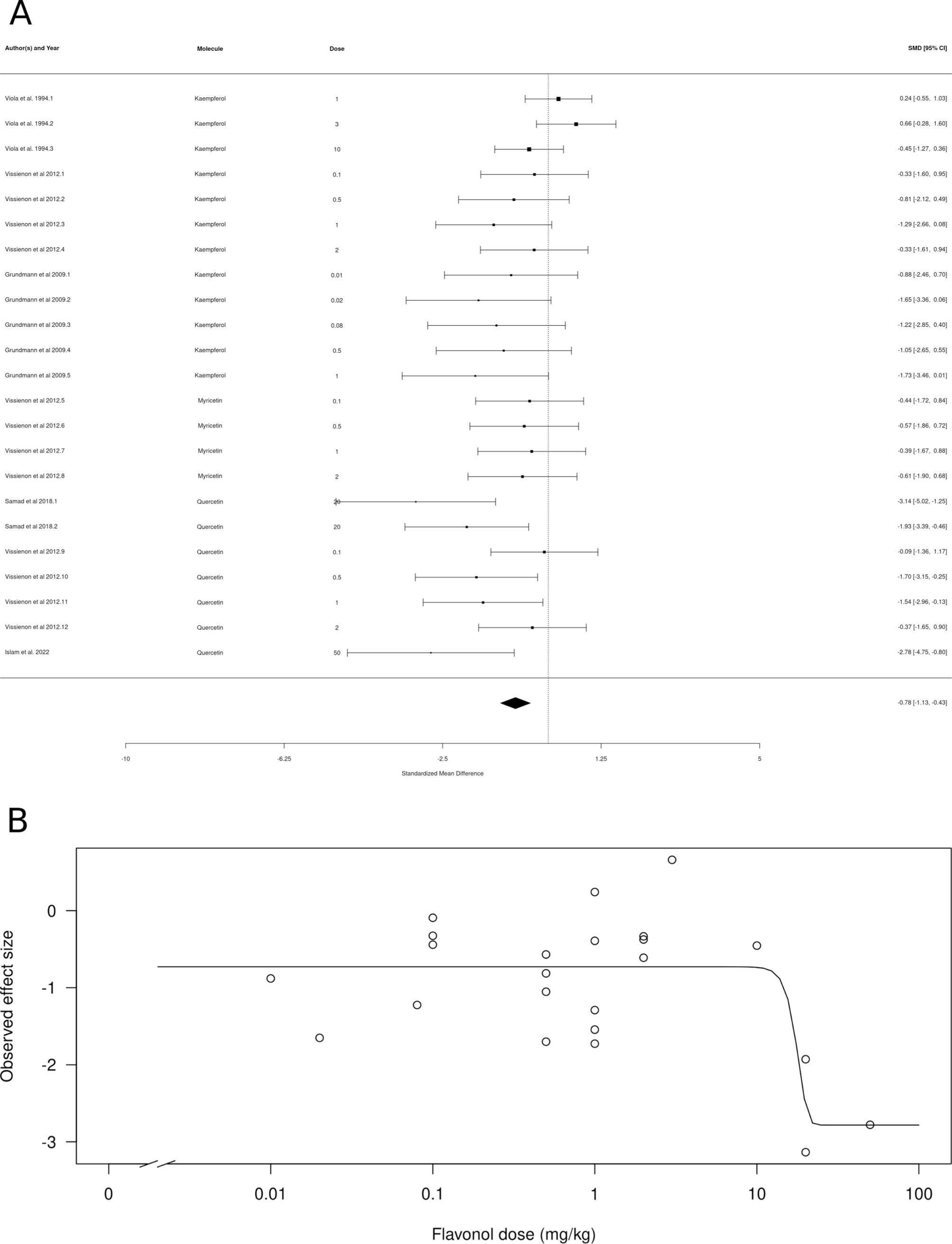
Subgroup analysis for flavanones. (A) Forest plot showing the results of 20 comparisons examining the effect of a flavanone on anxiety-like behavior in animal tests. The figure shows the standardized mean difference (SMD) between control and flavanone-treated groups with corresponding 95% confidence intervals in the individual comparison, based on a random-effects model. A negative standardized mean difference (SMD) corresponds to decreased anxiety-like behavior, while a positive SMD corresponds to increased anxiety-like behavior after flavanone treatment. The overall effect size is denoted by the diamond symbol. (B) Estimated dose-response curve, based on the observed effect size (SMD) with increasing flavanone dose. Each point corresponds to a single comparison from the forest plot. The curve was obtained by fitting generalized log-logistic 5 parameter models, with dose as independent variable and observed effect size SMD as dependent variable.

Flavanols (*k* = 4, represented only by (-)-epigallocatechin gallate) also appeared to produce an anxiolytic-like effect (Figure 8A). The estimated average standardized mean difference based on the random-effects model was 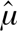 = -1.35 (95% CI: -2.07 to -0.63), interpreted as a medium-to-large effect size. Therefore, the average outcome differed significantly from zero (z = -3.6756, p = 0.0002), suggesting an overall anxiolytic-like effect. According to the Q-test, the true outcomes appear to be highly homogeneous (Q_[df= 3]_ = 0.3665, p = 0.912, τ² = 0, I² = 0.0%), as expected by an analysis that includes a single study. There is some evidence for a dose-dependent hormetic effect, with lower and higher doses producing slightly larger effects (Figure 8B), and a calculated ED _50_ of 75.56 ± 0.11 mg/kg (mean ± SEM).

**Figure 8.**
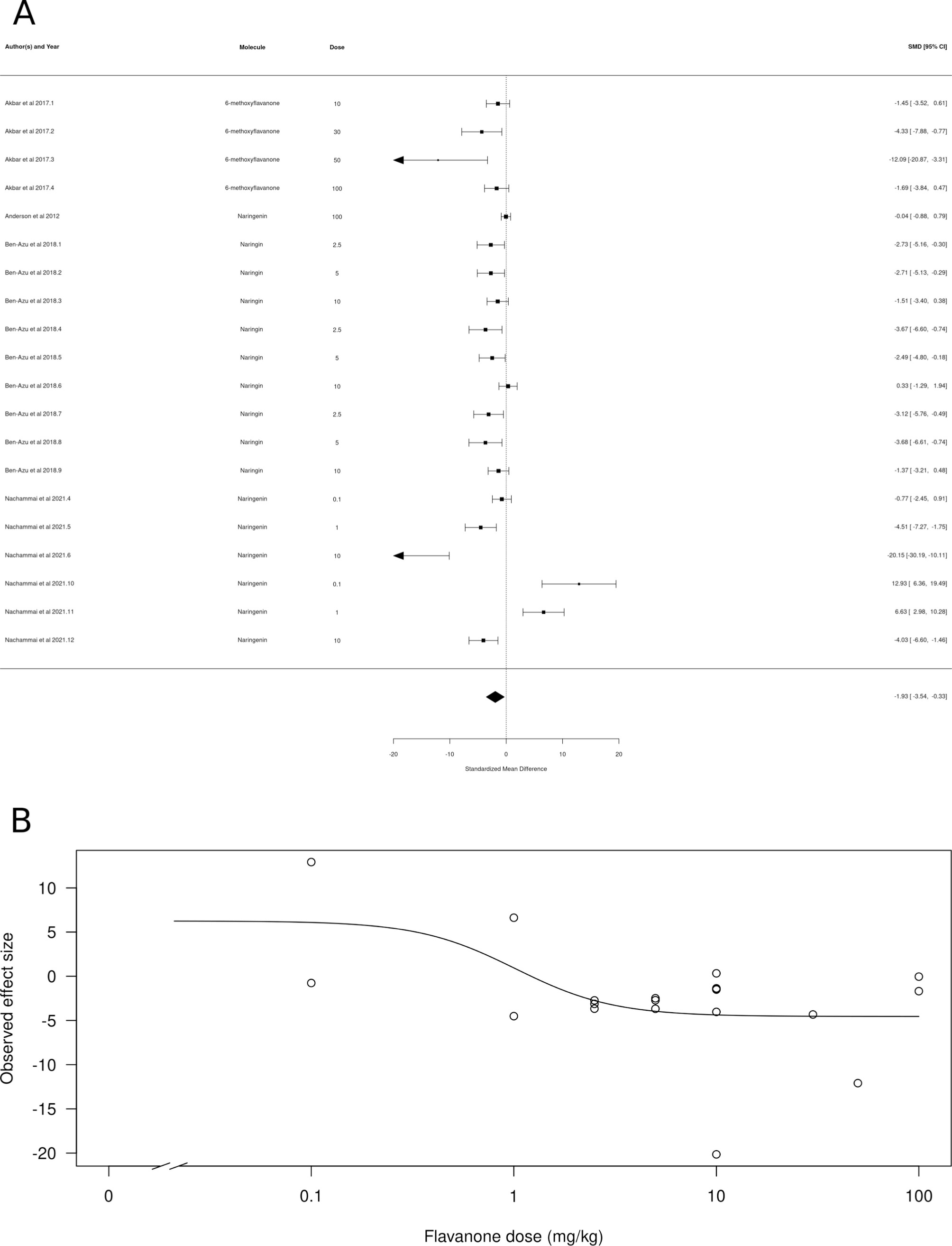
Subgroup analysis for flavanols. (A) Forest plot showing the results of 4 comparisons examining the effect of a flavanol on anxiety-like behavior in animal tests. The figure shows the standardized mean difference (SMD) between control and flavanol-treated groups with corresponding 95% confidence intervals in the individual comparison, based on a random-effects model. A negative standardized mean difference (SMD) corresponds to decreased anxiety-like behavior, while a positive SMD corresponds to increased anxiety-like behavior after flavanol treatment. The overall effect size is denoted by the diamond symbol. (B) Estimated dose-response curve, based on the observed effect size (SMD) with increasing flavanol dose. Each point corresponds to a single comparison from the forest plot. The curve was obtained by fitting a 5-parameter Brain-Cousens model, with dose as independent variable and observed effect size SMD as dependent variable.

Isoflavones (*k* = 3, represented only by puerarin) did not appear to produce anxiolytic- or anxiogenic-like effects (Figure 9A). The estimated average standardized mean difference based on the random-effects model was 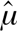 = -0.3065 (95% CI: -0.98 to 0.36). Therefore, the average outcome did not significantly differ from zero (z = -0.8978, p = 0.3693). In concordance with the wide range of SMDs, the true outcomes appear to be highly heterogeneous (Q_[df= 20]_ = 78.746, p < 0.0001, τ² = 1.889, I² = 83.17%). There is weak evidence for a dose-dependent effect, with higher doses producing slightly larger effects (Figure 9B), but it was not possible to calculate an ED_50_.

**Figure 9.**
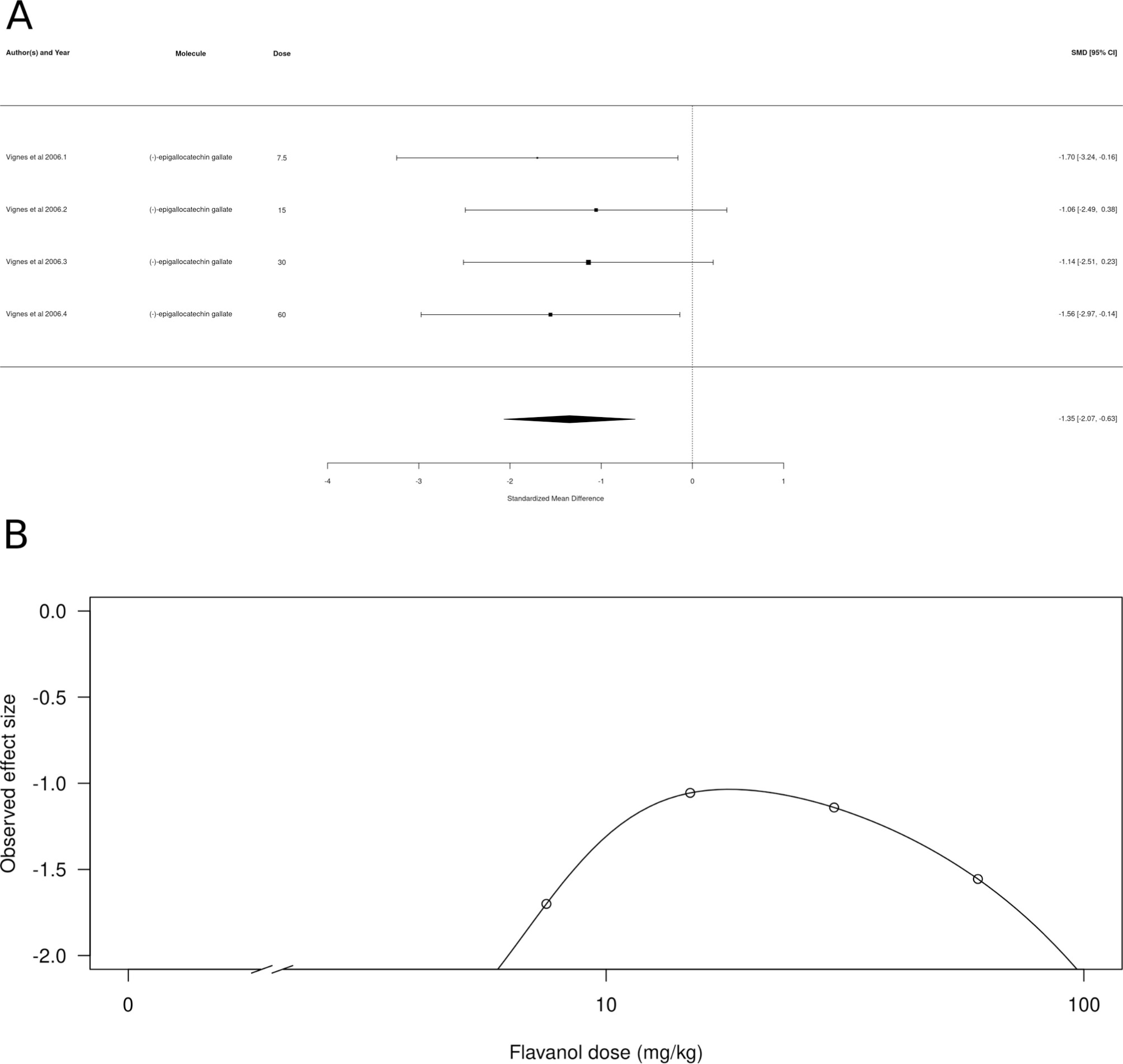
Subgroup analysis for isoflavones. (A) Forest plot showing the results of 3 comparisons examining the effect of a isoflavone on anxiety-like behavior in animal tests. The figure shows the standardized mean difference (SMD) between control and isoflavone-treated groups with corresponding 95% confidence intervals in the individual comparison, based on a random-effects model. A negative standardized mean difference (SMD) corresponds to decreased anxiety-like behavior, while a positive SMD corresponds to increased anxiety-like behavior after isoflavone treatment. The overall effect size is denoted by the diamond symbol. (B) Estimated dose-response curve, based on the observed effect size (SMD) with increasing isoflavone dose. Each point corresponds to a single comparison from the forest plot. The curve was obtained by fitting generalized log-logistic 5 parameter models, with dose as independent variable and observed effect size SMD as dependent variable.

Glycoside derivatives (*k* = 21) also did not appear to produce anxiolytic- or anxiogenic-like effects (Figure 10A). The estimated average standardized mean difference based on the random-effects model was 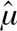 = -0.139 (95% CI: -0.86 to 0.58). Therefore, the average outcome did not significantly differ from zero (z = -0.3788, p = 0.7048). According to the Q-test, the true outcomes appear to be highly homogeneous (Q_[df= 2]_ = 0.2552, p = 0.8802, τ² = 0, I² = 0.0%), again as expected by an analysis that includes a single study. There is weak evidence for a dose-dependent effect, with higher doses producing larger effects (Figure 10B), with an ED_50_ of 4.93 ± 0.28 mg/kg (mean ± SEM).

**Figure 10.** Subgroup analysis for glycoside derivatives. (A) Forest plot showing the results of 21 comparisons examining the effect of a glycoside derivative on anxiety-like behavior in animal tests. The figure shows the standardized mean difference (SMD) between control and glycoside derivative-treated groups with corresponding 95% confidence intervals in the individual comparison, based on a random-effects model. A negative standardized mean difference (SMD) corresponds to decreased anxiety-like behavior, while a positive SMD corresponds to increased anxiety-like behavior after glycoside derivative treatment. The overall effect size is denoted by the diamond symbol. (B) Estimated dose-response curve, based on the observed effect size (SMD) with increasing glycoside derivative dose. Each point corresponds to a single comparison from the forest plot. The curve was obtained by fitting generalized log-logistic 5 parameter models, with dose as independent variable and observed effect size SMD as dependent variable.

Glucoside derivatives (*k* = 3, represented only by apigenin 7-glucoside) appeared to produce an anxiolytic-like effect (Figure 11A). The estimated average standardized mean difference based on the random-effects model was 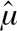 = -2.29 (95% CI: -3.57 to -1.01), interpreted as a large effect size. Therefore, the average outcome differed significantly from zero (z = -3.4919, p = 0.0005), suggesting an overall anxiolytic-like effect. According to the Q-test, the true outcomes appear to be highly homogeneous (Q_[df= 2]_ = 0.2525, p = 0.8814, τ² = 0, I² = 0.0%), as expected by an analysis that includes a single study. There is some evidence for a dose-dependent effect, with lower doses producing larger effects (Figure 11B), albeit with high variability: a calculated ED_50_ of 3.84 ± 356.94 mg/kg (mean ± SEM). 687.98 mg/kg (mean ± SEM).

**Figure 11.**
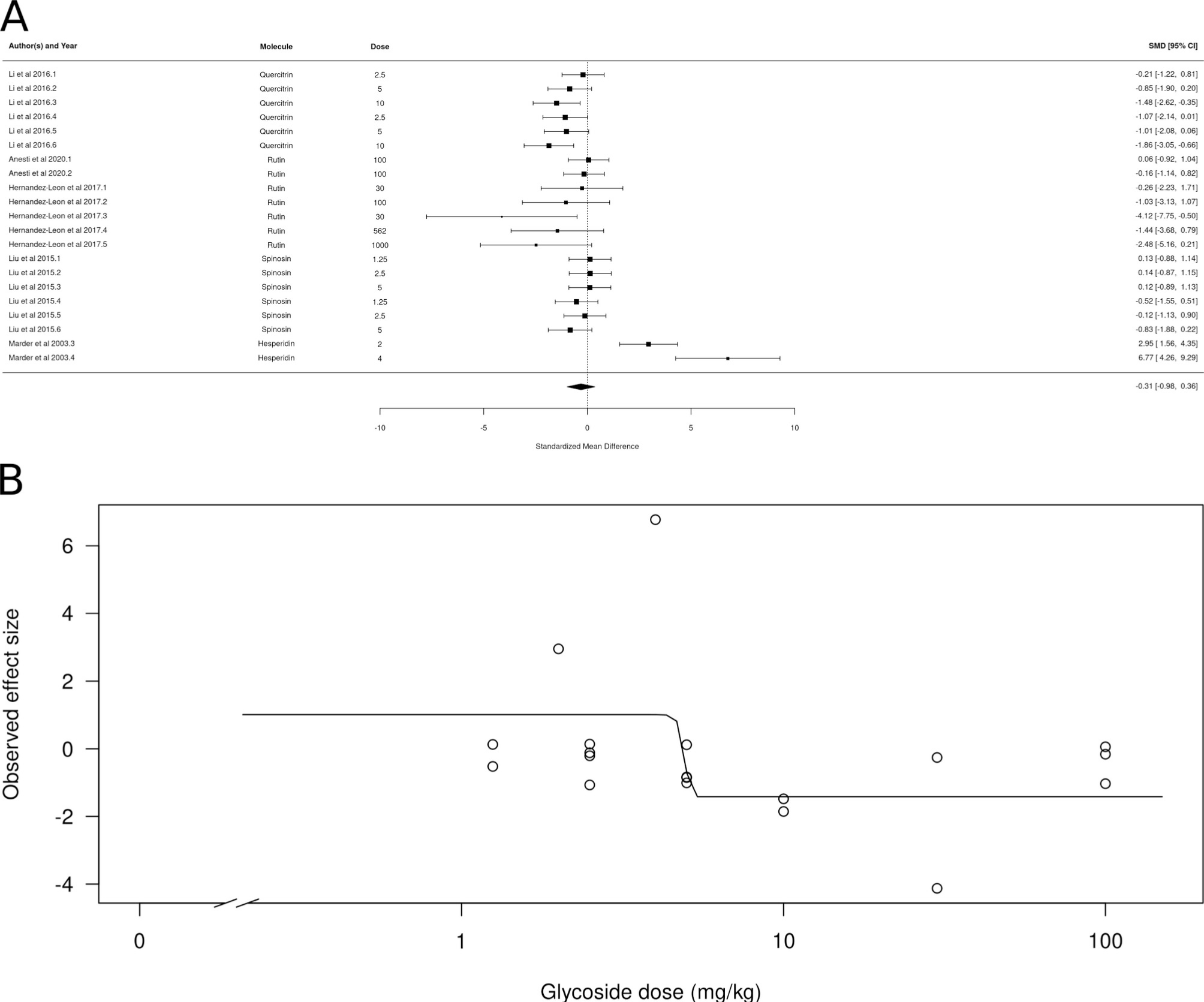
Subgroup analysis for glucoside derivatives. (A) Forest plot showing the results of 3 comparisons examining the effect of a glucoside derivative on anxiety-like behavior in animal tests. The figure shows the standardized mean difference (SMD) between control and glucoside derivative-treated groups with corresponding 95% confidence intervals in the individual comparison, based on a random-effects model. A negative standardized mean difference (SMD) corresponds to decreased anxiety-like behavior, while a positive SMD corresponds to increased anxiety-like behavior after glucoside derivative treatment. The overall effect size is denoted by the diamond symbol. (B) Estimated dose-response curve, based on the observed effect size (SMD) with increasing glucoside derivative dose. Each point corresponds to a single comparison from the forest plot. The curve was obtained by fitting generalized log-logistic 5 parameter models, with dose as independent variable and observed effect size SMD as dependent variable.

Finally, flavonolignans (*k* = 6, represented only by silibinin) did not appear to produce neither anxiolytic-like nor anxiogenic-like effects (Figure 12A). The estimated average standardized mean difference based on the random-effects model was *μ* = -4.86 (95% CI: -14.51 to 4.79). Therefore, the average outcome did not significantly differ from zero (z = -0.9867, p = 0.3238). According to the Q-test, the true outcomes appear to be highly heterogeneous (Q_[df= 5]_ = 81.86, p < 0.0001, τ² = 135.7513, I² = 96.8%). There is some evidence for a dose-dependent effect, with higher doses producing larger effects (Figure 12B), albeit with high variability: a calculated ED_50_ of 4.04 ±

**Figure 12.**
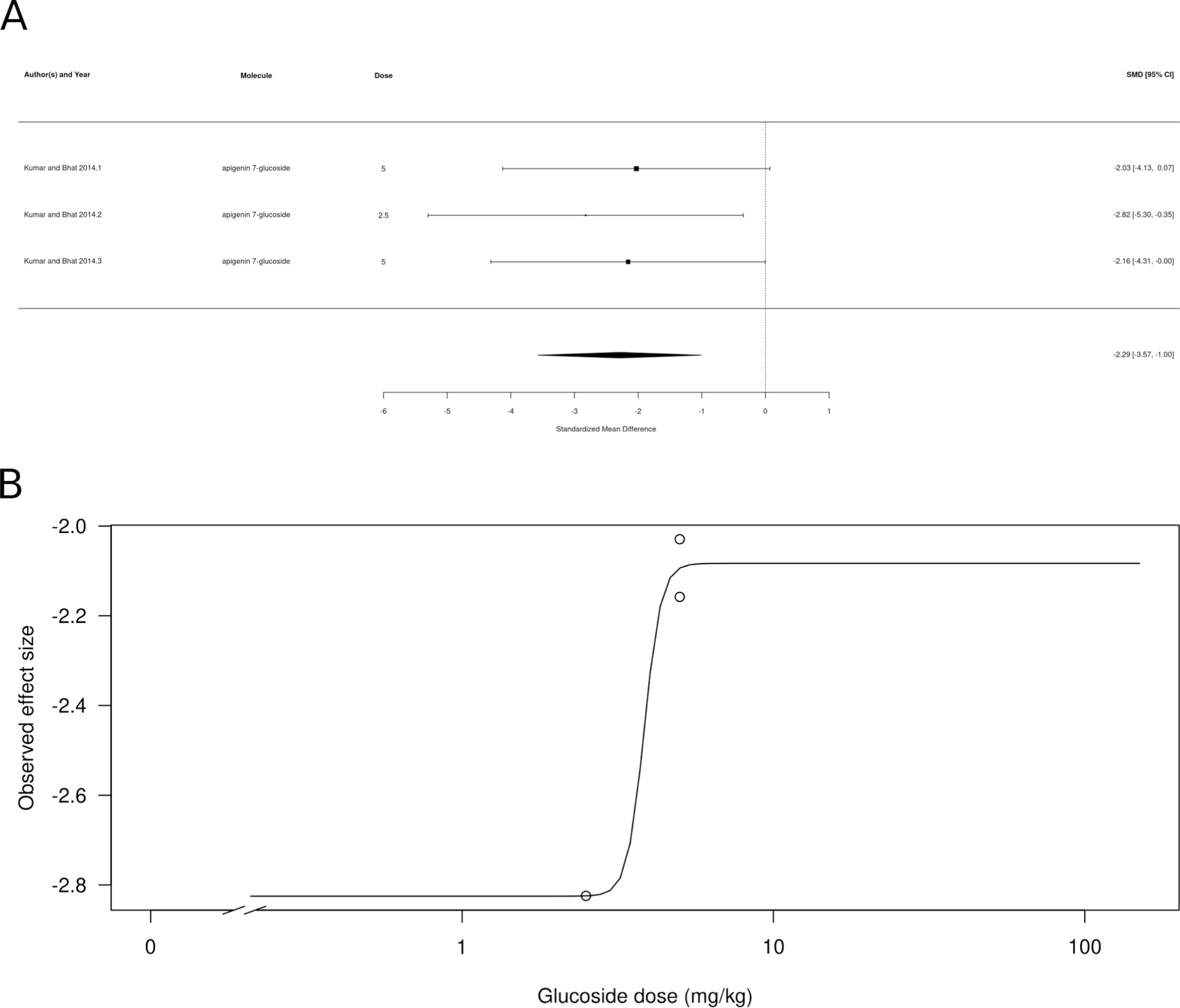
Subgroup analysis for flavonolignans. (A) Forest plot showing the results of 6 comparisons examining the effect of a flavonolignan on anxiety-like behavior in animal tests. The figure shows the standardized mean difference (SMD) between control and flavonolignan-treated groups with corresponding 95% confidence intervals in the individual comparison, based on a random-effects model. A negative standardized mean difference (SMD) corresponds to decreased anxiety-like behavior, while a positive SMD corresponds to increased anxiety-like behavior after flavonolignan treatment. The overall effect size is denoted by the diamond symbol. (B) Estimated dose-response curve, based on the observed effect size (SMD) with increasing flavonolignan dose. Each point corresponds to a single comparison from the forest plot. The curve was obtained by fitting generalized log-logistic 5 parameter models, with dose as independent variable and observed effect size SMD as dependent variable.

## 4. Discussion

The present work presents bibliometric analysis on the literature on the acute and chronic effects of natural and synthetic flavonoids on anxiety-like behavior in animal tests, followed by a meta-analysis of filtered data. Bibliometric analysis suggested a prolific field that can improve in terms of international collaboration; thematic analysis suggests that the field was highly focused on testing drug effects, with part of it focusing on mechanisms – especially neurobiological effects on oxidative stress and neuropathology –, with few emerging themes. The meta-analysis revealed a general anxiolytic-like effect was found for these molecules, with a high heterogeneity in findings. Subgroup analyses found that acute treatment produced significant effects, while no evidence for chronic effects were found. Moreover, it was found that, for all molecule classes included in the study, only isoflavones, glycoside derivatives, and flavanolignans did not show evidence of an anxiolytic-like effect. Finally, we found evidence for publication bias, suggesting a “file-drawer effect”.

### 4.1. Mapping the field with bibliometric analyses

Bibliometric analysis suggest that the field is highly concentrated on few research groups that are mostly located in the Global South, including Asian, African, Middle Eastern, and Latin American countries. These results are not surprising, considering that China and India are the top two research producing countries in the field of flavonoids, both in plant physiology (Y. Li et al., 2023) and health (Perez-Vizcaino & Fraga, 2018). Li et al. (2023) attribute part of this trend on the rapid growth of agricultural research and production in these countries. International collaborations are still very incipient, with most research groups being formed by individuals from the same institution, and a low proportion multiple country publications. Improving international collaborations seems to be an important step forward in the field, but the current framework for international collaboration policies still implies the construction of inequalities in capabilities and abilities, with lack of reciprocity and agency (Martinez & Sá, 2020; Pineda et al., 2020; Skupien & Rüffin, 2020). Considering that most of the research on anxiolytic-like effects of flavonoids tend to happen on the “Global South”, strategies to mitigate these inequalities are needed.

The analysis of trending topics suggest that the field is still very much dominated by exploratory research on putative behavioral effects of flavonoids and its phytochemical characterization. This is suggested by the fact that the “motor themes” (those with high centrality and density in a given field, being both well developed and important for the structure of the area) involve mainly areas of phytochemistry, including compound extraction and quantification methods. Moreover, the “core sources” for research on anxiolytic-like effects of flavonoids include mainly journals from the area of pharmacology. Conversely, the basic themes (which involve themes that are important for a given research field, but are underdeveloped) include basic techniques in behavioral pharmacology (dose-response relationships, different behavioral methods), as well as general mechanisms of drug effect. This is in contrast with the multitude of known mechanisms of action of flavonoids, which include modulation of GABAergic receptors (Hanrahan et al., 2011; Wasowski & Marder, 2012), monoamine metabolism (Carradori et al., 2016), and antioxidant properties (Hritcu et al., 2017). Niche and emerging or declining themes, which show either specialized or peripheral themes with low external ties to other themes, included the study of inflammation and neuropathology mechanisms, as well as the use of computational techniques. This suggests that the field, as a whole, could benefit from more mechanistic and confirmatory research.

### 4.2. Systematic review and meta-analysis

In contrast with the bibliometric analysis (which included all documents from the area retrieved from PubMed, not only those which met the inclusion criteria for the meta-analysis), a systematic review of the experimental literature suggested a field that is more concerned with mechanisms. Of the 34 documents retained for the analysis, the majority also studied mechanisms of action, either indirectly (e.g., with competitive radioligand binding assays, or *in silico* molecular docking studies) or directly, by pre-treating animals with different receptor antagonists. Most of these studies focused on GABA_A_ receptors, with analysis of either the orthosteric site for GABA or different allosteric sites. A few (e.g., Ferreira et al., 2020; J. Liu et al., 2015) also investigated the participation of the serotonergic system.

The general meta-analysis found evidence for an anxiolytic-like effect in preclinical animal tests, albeit with high heterogeneity. The overall effect size is larger than that reported for clinical studies with flavonoids (Jia et al., 2023) or BZDs (Gomez et al., 2018) for the treatment of anxiety; this is probably due to the much higher heterogeneity in clinical studies that, by nature, lack experimental control. The high heterogeneity found in the general meta-analysis prompted us to investigate its sources, focusing on treatment duration and molecule class.

Differences in the effects of treatment duration are highly relevant from a translational point of view, with implications for treatment regimens in humans. While BZDs are commonly prescribed for acute and subacute use, they produce tolerance (Shinfuku et al., 2019), and important adverse effects (L. Liu et al., 2020) and risk of addiction (Longo & Johnson, 2000) increase with chronic use. Nonetheless, a meta-analysis of clinical studies with flavonoids suggested that chronic treatment appears to produce better results (Jia et al., 2023). However, the fact that we found anxiolytic-like effects of acute, but not chronic, treatment can also be interpreted in terms of number of studies, since only 4 of the 34 studies included in the meta-analysis attempted chronic treatment. Further preclinical studies using flavonoids should incorporate chronic treatment, testing not only for the development of tolerance, but also the development of adverse effects, withdrawal-like symptoms, and abuse potential.

We also found molecule class-dependent anxiolytic-like effects in the meta-analysis. Subgroup analysis found significant effects of flavones, chalcones, flavonols, flavanones, flavanols, and glycoside derivatives. The majority of studies involved flavones and chalcone, which also had the larger effect sizes. 2′-methoxy-6-methylflavone and 3-hydroxy-2’-methoxy-6-methylflavone modulate the activity of GABAA receptors in subunit-selective ways: the first appears to directly activate α2/γ2-containing GABAA receptors, while the second acts as a positive allosteric modulation of the α2β2/3γ2L and direct activation of α4β2/3δ GABAA receptors (Karim et al., 2011, 2012). 5-methoxyflavone also appears to bind α2-containing receptors (Shanmugasundaram et al., 2020). Chrysin, wogonin, 6-methylapigenin, luteonin, and 5,7,2′-trihydroxy-6,8-dimethoxyflavone all appear to have affinity for the central BZD binding site (Wasowski & Marder, 2012). Apigenin, on the other hand, appears to act on GABA_A_ receptors either as an inverse agonist or as an antagonist at the central BZD binding site (Avallone et al., 2000; Dekermendjian et al., 1999; Zanoli et al., 2000), and inspection of the forest plot in Figure 4A suggest that this molecule does not have a clear anxiolytic-like effect in animal tests. There is also some evidence for a participation of the central BZD site on the effects of chalcones, at least in zebrafish (Ferreira et al., 2020, 2021; Santos Oliveira et al., 2024). On the other hand, flavones and chalcones also appear to act at other targets, including monoaminergic neurotransmission (Carradori et al., 2016) and redox balance (Hritcu et al., 2017). With few exceptions, most of the work on the effects of flavones and chalcones on anxiety-like behavior did not explicitly test the hypothesis of a mediation by either of these systems.

Interestingly, in the case of chalcones, the dose-response curve from the meta-analysis suggested an hormetic effect, with lower and higher doses producing a more prominent anxiolytic-like effect. The flavanones naringenin, naringin, and 6-methoxyflavanone; the flavanol (-)-epigallocatechin gallate; and the glucoside derivative apigenin 7-glucoside all appeared to decrease anxiety-like behavior. (-)-epigallocatechin gallate appears to produce an hormetic dose-response profile, and the effects of apigenin 7-glucoside appear to decrease with increasing dose. Hormesis is defined as a biphasic response, with stimulatory and inhibitory responses that can be either be mediated by a single receptor recruiting different downstream signaling mechanisms, by different receptors to which the molecule has different affinities, or by different targets at different levels of a signaling pathway (Calabrese, 2013; Calabrese & Baldwin, 2002). There is some evidence for both GABAergic and serotonergic mechanisms in mediating the behavioral effects of anxiolytic chalcones (Ferreira et al., 2019, 2020; Mendes et al., 2023); however, since assessing hormetic mechanisms implicates blocking receptors at both the inhibitory and stimulatory doses of the chalcone (Calabrese, 2013), currently there is no evidence supporting any specific hormetic mechanism for these molecules.

The flavonols kaempferol, quercetin, and myricetin also showed evidence for an anxiolytic-like effect. The mechanisms of action of these flavanols are uncertain; intracerebroventricular injection of kaempferol reduces anxiety-like behavior in male rats by a GABA_A_ receptor-dependent mechanism (Zarei et al., 2021). However, intraperitoneal injections of these three flavanols failed to affect anxiety-like behavior, while administration *per os* decreased it, suggesting that these flavanols are pro-drugs which are metabolized before they reach the brain (Vissiennon et al., 2012).

Interestingly, no effects were found for isoflavones, a class of molecules that is also classified as a phytoestrogen. The isoflavone content of the diet has been associated with increased anxiety-like behavior in male rats (Hartley et al., 2003), but in non-gonadotomized female rats isoflavone supplementation has an anxiolytic-like effect (Friedman & Frye, 2011). In a meta-analysis of the outcomes of dietary supplementation of flavonoids in women, isoflavones failed to produce an effect on anxiety outcomes (Jia et al., 2023). Again, care must be taken in the extrapolation of the results from the present meta-analysis, given that only one study using a single isoflavone, puerarin, was included in the analysis, and the study used only acute treatment.

### 4.3. Recommendations for future research

Overall, the effects of flavonoids in preclinical behavioral research appear to indicate a class-dependent effect, more evident at acute than chronic treatments, that are hypothesized to be mediated by GABAergic and/or 5-HTergic mechanisms, although hypothesis testing of mechanisms is still weak. As a result, there are still many open questions in preclinical research of the putative anxiolytic effects of flavonoids to be answered before these molecules are advanced in the drug development pipeline. Most of the outstanding questions are related to basic pharmacological and psychopharmacological research. Based on the bibliometric analysis, as well as on the systematic review and meta-analysis, we recommend that:

1. **The field needs to further explore mechanistic hypothesis-driven research**. While the in the initial stages of drug development exploratory research (including the establishment of dose-response curves, ligand binding and/or molecular docking assays, and the use of behavioral test batteris) is crucial, once these are established the search for mechanisms is fundamental to establish a good evidence base (Crabbe & Morris, 2004; Stegenga, 2022).
2. **In order to do so, increasing collaboration between groups could be important.** Mechanistic research usually involves the use of multiple techniques, from ligand binding, molecular docking, and electrophysiology and/or signal transduction biochemistry, to molecular biology and behavioral pharmacology assays. Not all research groups will be able to tackle all the questions that are important for mechanistic research – especially considering that most of the groups currently working in the field come from the Global South. Thus, increasing collaboration between research groups could be important to move the field towards more mechanistic, hypothesis-driven research.
3. **Improving reproducibility and increasing power is paramount**. Most of the documents found in the meta-analysis rely on very small sample sizes, typical of exploratory research, which can lead to inflated effect sizes and might be responsible for the high heterogeneity. As research in the field advances, sample sizes will need to be increased in order to approach true effects and reduce heterogeneity. Moreover, steps towards increasing reproducibility are also direly needed (Begley & Ioannidis, 2015; Richter et al., 2009); while the present work did not assess risk of bias, a cursory view of the Methods section of most documents included in the meta-analysis reveals that it is impossible to discern whether or not blinding or randomization was applied. Again, multi-site collaborations could help, as introducing heterogeneity in these studies (with each laboratory covering a portion of the final sample size) appears to increase reproducibility while decreasing the need for a single group bearing the weight of large sample sizes (Drude et al., 2022; Voelkl et al., 2018).

Flavonoids can represent a very promising lead in anxiolytic research, a field that is in dire need of novel molecules that can be translated from preclinical to clinical research. However, in order for that potential to be realized, research practices in the field need to advance. We hope that, in the near future, the field of flavonoid research reaches the maturity it needs.

## Acknowledgments

This work was supported by a grant from Conselho Nacional de Desenvolvimento Científico e Tecnológico (CNPq/Brazil) grant to CM (grant #406336/2023-7).

---

**Figure S1.**
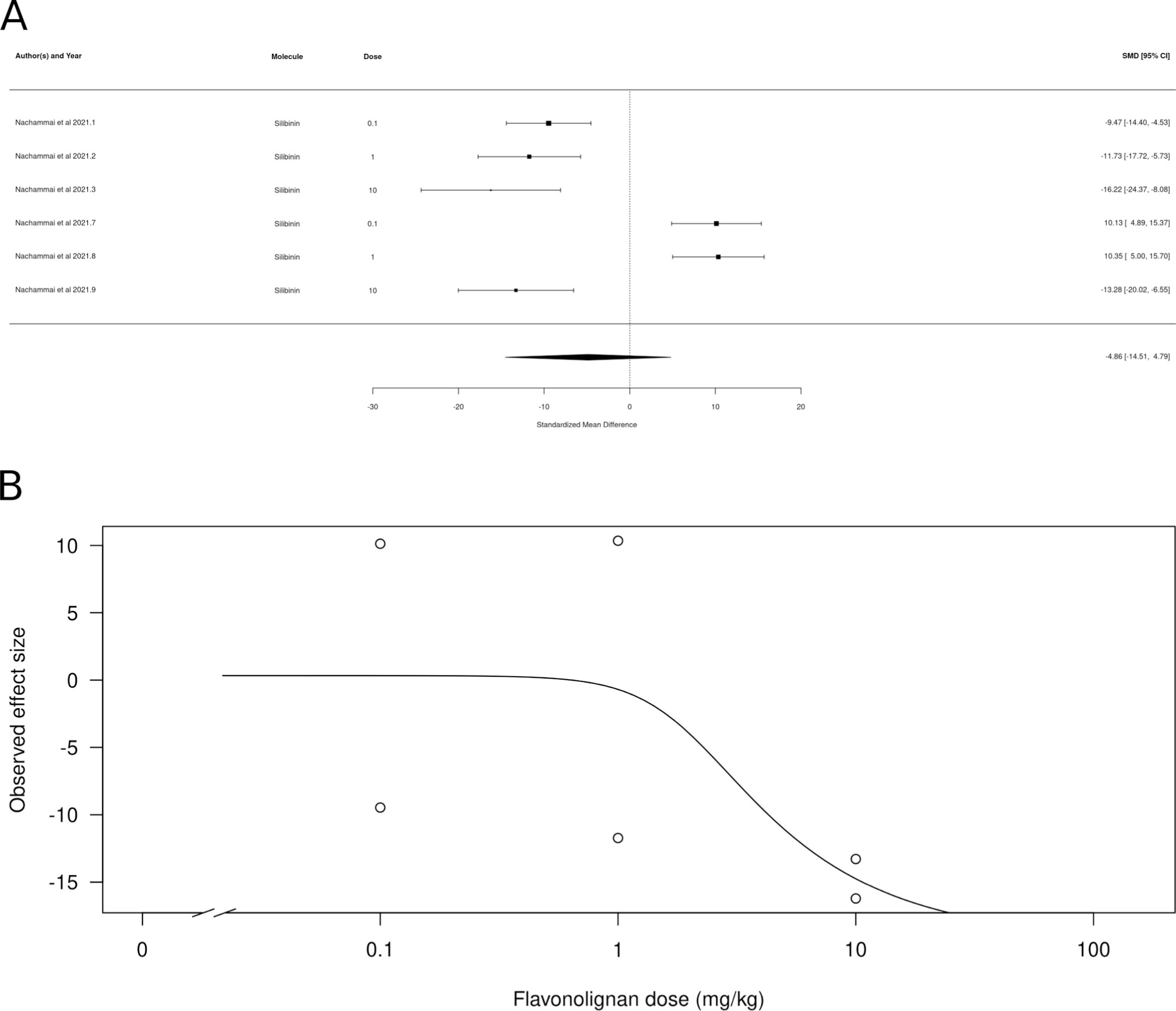
Forest plot showing the results of the overall meta-analysis, with 172 comparisons examining the effect of a flavonoid on anxiety-like behavior in animal tests. The figure shows the standardized mean difference (SMD) between control and flavonoid-treated groups with corresponding 95% confidence intervals in the individual comparison, based on a random-effects model. A negative standardized mean difference (SMD) corresponds to decreased anxiety-like behavior, while a positive SMD corresponds to increased anxiety-like behavior after flavonoid treatment. The overall effect size is denoted by the diamond symbol.

Figure S2 Forest plot showing the results of the subgroup meta-analysis for acute treatment, with 150 comparisons examining the effect of a flavonoid on anxiety-like behavior in animal tests in acute treatment. The figure shows the standardized mean difference (SMD) between control and flavonoid-treated groups with corresponding 95% confidence intervals in the individual comparison, based on a random-effects model. A negative standardized mean difference (SMD) corresponds to decreased anxiety-like behavior, while a positive SMD corresponds to increased anxiety-like behavior after flavonoid treatment. The overall effect size is denoted by the diamond symbol.

Figure S3 Forest plot showing the results of the subgroup meta-analysis for chronic treatment, with 22 comparisons examining the effect of a flavonoid on anxiety-like behavior in animal tests in chronic treatment. The figure shows the standardized mean difference (SMD) between control and flavonoid-treated groups with corresponding 95% confidence intervals in the individual comparison, based on a random-effects model. A negative standardized mean difference (SMD) corresponds to decreased anxiety-like behavior, while a positive SMD corresponds to increased anxiety-like behavior after flavonoid treatment. The overall effect size is denoted by the diamond symbol.

## References

Abou Baker, D. H. (2022). An ethnopharmacological review on the therapeutical properties of flavonoids and their mechanisms of actions: A comprehensive review based on up to date knowledge. Toxicology Reports, 9, 445–469. 10.1016/j.toxrep.2022.03.011

Akbar, S., Subhan, F., Karim, N., Aman, U., Ullah, S., Shahid, M., Ahmad, N., Fawad, K., & Sewell, R. D. E. (2017). Characterization of 6-methoxyflavanone as a novel anxiolytic agent: A behavioral and pharmacokinetic approach. European Journal of Pharmacology, 801, 19–27. 10.1016/j.ejphar.2017.02.047

Alkin, B. (2017). Useful plants—Medicines. Em K. J. Willis (Org.), State of the World’s Plants 2017. Royal Botanic Gardens, Kew. https://www.ncbi.nlm.nih.gov/books/NBK464488/

Amin, F., Ibrahim, M. A. A., Rizwan-ul-Hasan, S., Khaliq, S., Gabr, G. A., Muhammad, Khan, A., Sidhom, P. A., Tikmani, P., Shawky, A. M., Ahmad, S., & Abidi, S. H. (2022). Interactions of Apigenin and Safranal with the 5HT1A and 5HT2A Receptors and Behavioral Effects in Depression and Anxiety: A Molecular Docking, Lipid-Mediated Molecular Dynamics, and In Vivo Analysis. Molecules, 27, Artigo 24. 10.3390/molecules27248658

Anderson, W., Barrows, M., Lopez, F., Rogers, S., Ortiz-Coffie, A., Norman, D., Hodges, J., McDonald, K., Barnes, D., McCall, S., Don, J. A., & Ceremuga, T. E. (2012). Investigation of the anxiolytic effects of naringenin, a component of *Mentha aquatica*, in the male Sprague-Dawley rat. Holistic Nursing Practice, 26, 52–57. 10.1097/HNP.0b013e31823c003a

Anesti, M., Stavropoulou, N., Atsopardi, K., Lamari, F. N., Panagopoulos, N. T., & Margarity, M. (2020). Effect of rutin on anxiety-like behavior and activity of acetylcholinesterase isoforms in specific brain regions of pentylenetetrazol-treated mice. Epilepsy & Behavior, 102, 106632. 10.1016/j.yebeh.2019.106632

Antunes, M. S., Cattelan Souza, L., Ladd, F. V. L., Ladd, A. A. B. L., Moreira, A. L., Bortolotto, V. C., Silva, M. R. P., Araújo, S. M., Prigol, M., Nogueira, C. W., & Boeira, S. P. (2020). Hesperidin Ameliorates Anxiety-Depressive-Like Behavior in 6-OHDA Model of Parkinson’s Disease by Regulating Striatal Cytokine and Neurotrophic Factors Levels and Dopaminergic Innervation Loss in the Striatum of Mice. Molecular Neurobiology, 57, 3027–3041. 10.1007/s12035-020-01940-3

Aria, M., & Cuccurullo, C. (2017). *bibliometrix*: An R-tool for comprehensive science mapping analysis. Journal of Informetrics, 11(4), 959–975. 10.1016/j.joi.2017.08.007

Avallone, R., Zanoli, P., Puia, G., Kleinschnitz, M., Schreier, P., & Baraldi, M. (2000). Pharmacological profile of apigenin, a flavonoid isolated from *Matricaria chamomilla*. Biochemical Pharmacology, 59, 1387–1394. 10.1016/s0006-2952(00)00264-1

Begg, C. B., & Mazumdar, M. (1994). Operating Characteristics of a Rank Correlation Test for Publication Bias. Biometrics, 50, 1088–1101. 10.2307/2533446

Begley, C. G., & Ioannidis, J. P. A. (2015). Reproducibility in Science. Circulation Research, 116(1), 116–126. 10.1161/CIRCRESAHA.114.303819

Ben-Azu, B., Nwoke, E. E., Umukoro, S., Aderibigbe, A. O., Ajayi, A. M., & Iwalewa, E. O. (2018). Evaluation of the Neurobehavioral Properties of Naringin in Swiss Mice. Drug Research, 68, 465–474. 10.1055/a-0575-3730

Bradford, S. C. (1985). Sources of information on specific subjects. Journal of Information Science, 10, 176–180. 10.1177/016555158501000407 (Trabalho original publicado 1934)

Brain, P., & Cousens, R. (1989). An equation to describe dose responses where there is stimulation of growth at low doses. Weed Research, 29, 93–96. 10.1111/j.1365-3180.1989.tb00845.x

Calabrese, E. J. (2013). Hormetic mechanisms. Critical Reviews in Toxicology, 43, 580–606. 10.3109/10408444.2013.808172

Calabrese, E. J., & Baldwin, L. A. (2002). Defining hormesis. Human & Experimental Toxicology, 21, 91–97. 10.1191/0960327102ht217oa

Carradori, S., Gidaro, M. C., Petzer, A., Costa, G., Guglielmi, P., Chimenti, P., Alcaro, S., & Petzer, J. P. (2016). Inhibition of Human Monoamine Oxidase: Biological and Molecular Modeling Studies on Selected Natural Flavonoids. Journal of Agricultural and Food Chemistry, 64, 9004–9011. 10.1021/acs.jafc.6b03529

Cochran, W. G. (1954). The combination of estimates from different experiments. Biometrics, 10, 101–129. 10.2307/3001666

Cohen, J. (1988). *Statistical power analysis for the behavioral sciences* (2nd ed). Routledge.

Coleta, M., Campos, M. G., Cotrim, M. D., Lima, T. C. M. de, & Cunha, A. P. da. (2008). Assessment of luteolin (3’,4’,5,7-tetrahydroxyflavone) neuropharmacological activity. Behavioural Brain Research, 189, 75–82. 10.1016/j.bbr.2007.12.010

Crabbe, J. C., & Morris, R. G. M. (2004). Festina lente: Late-night thoughts on high-throughput screening of mouse behavior. Nature Neuroscience, 7, 1175–1179. 10.1038/nn1343

Cueto-Escobedo, J., Andrade-Soto, J., Lima-Maximino, M., Maximino, C., Hernández-López, F., & Rodríguez-Landa, J. F. (2020). Involvement of GABAergic system in the antidepressant-like effects of chrysin (5,7-dihydroxyflavone) in ovariectomized rats in the forced swim test: Comparison with neurosteroids. Behavioural Brain Research, 386, 112590. 10.1016/j.bbr.2020.112590

Cueto-Escobedo, J., García-García, F., Guillén-Ruiz, G., & Rodríguez-Landa, J. F. (2020). Anxiolytic effects of fluoxetine in preclinical and clinical research. Em L. V. Berhardt (Org.), Advances in medicine and biology (p. 104–137). Nova Science Publisher, Inc.

Cueto-Escobedo, J., German-Ponciano, L. J., Guillén-Ruiz, G., Soria-Fregozo, C., & Herrera-Huerta, E. V. (2022). Zebrafish as a Useful Tool in the Research of Natural Products With Potential Anxiolytic Effects. Frontiers in Behavioral Neuroscience, 15, 795285. 10.3389/fnbeh.2021.795285

da Silva Chaves, S. N., Felício, G. R., Costa, B. P. D., de Oliveira, W. E. A., Lima-Maximino, M. G., Siqueira Silva, D. H. de, & Maximino, C. (2018). Behavioral and biochemical effects of ethanol withdrawal in zebrafish. Pharmacology Biochemistry and Behavior, 169, 48–58. 10.1016/j.pbb.2018.04.006

Dekermendjian, K., Kahnberg, P., Witt, M. R., Sterner, O., Nielsen, M., & Liljefors, T. (1999). Structure-activity relationships and molecular modeling analysis of flavonoids binding to the benzodiazepine site of the rat brain GABA(A) receptor complex. Journal of Medicinal Chemistry, 42, 4343–4350. 10.1021/jm991010h

Drude, N. I., Martinez-Gamboa, L., Danziger, M., Collazo, A., Kniffert, S., Wiebach, J., Nilsonne, G., Konietschke, F., Piper, S. K., Pawel, S., Micheloud, C., Held, L., Frommlet, F., Segelcke, D., Pogatzki-Zahn, E. M., Voelkl, B., Friede, T., Brunner, E., Dempfle, A., … Toelch, U. (2022). Planning preclinical confirmatory multicenter trials to strengthen translation from basic to clinical research – a multi-stakeholder workshop report. Translational Medicine Communications, 7, 24. 10.1186/s41231-022-00130-8

Egger, M., Smith, G. D., Schneider, M., & Minder, C. (1997). Bias in meta-analysis detected by a simple, graphical test. BMJ, 315, 629–634. 10.1136/bmj.315.7109.629

Ferreira, M. K. A., da Silva, A. W., Dos Santos Moura, A. L., Sales, K. V. B., Marinho, E. M., do Nascimento Martins Cardoso, J., Marinho, M. M., Bandeira, P. N., Magalhães, F. E. A., Marinho, E. S., de Menezes, J. E. S. A., & Dos Santos, H. S. (2021). Chalcones reverse the anxiety and convulsive behavior of adult zebrafish. Epilepsy & Behavior, 117, 107881. 10.1016/j.yebeh.2021.107881

Ferreira, M. K. A., da Silva, A. W., Silva, F. C. O., Holanda, C. L. A., Barroso, S. M., Lima, J. D. R., Vieira Neto, A. E., Campos, A. R., Bandeira, P. N., Dos Santos, H. S., de Lemos, T. L. G., Siqueira, S. M. C., Magalhães, F. E. A., & de Menezes, J. E. S. A. (2019). Anxiolytic-like effect of chalcone N-{(4’-[(E)-3-(4-fluorophenyl)-1-(phenyl) prop-2-en-1-one]} acetamide on adult zebrafish (Danio rerio): Involvement of the GABAergic system. Behavioural Brain Research, 374, 111871. 10.1016/j.bbr.2019.03.040

Ferreira, M. K. A., da Silva, A. W., Silva, F. C. O., Vieira Neto, A. E., Campos, A. R., Alves Rodrigues Santos, S. A., Rodrigues Teixeira, A. M., da Cunha Xavier, J., Bandeira, P. N., Sampaio Nogueira, C. E., de Brito, D. H. A., Rebouças, E. L., Magalhães, F. E. A., de Menezes, J. E. S. A., & dos Santos, H. S. (2020). Anxiolytic-like effect of chalcone N-{4’[(*2E*)-3-(3-nitrophenyl)-1-(phenyl)prop-2-en-1-one]} acetamide on adult zebrafish (*Danio rerio*): Involvement of the 5-HT system. Biochemical and Biophysical Research Communications, 526(2), 505–511. 10.1016/j.bbrc.2020.03.129

Friedman, J., & Frye, C. (2011). Anti-anxiety, cognitive, and steroid biosynthetic effects of an isoflavone-based dietary supplement are gonad and sex-dependent in rats. Brain Research, 1379, 164–175. 10.1016/j.brainres.2010.12.025

Gencturk, S., & Unal, G. (2024). Rodent tests of depression and anxiety: Construct validity and translational relevance. *Cognitive, Affective*, & Behavioral Neuroscience, 24, 191–224. 10.3758/s13415-024-01171-2

Gerlai, R. (2023). Zebrafish (*Danio rerio*): A newcomer with great promise in behavioral neuroscience. Neuroscience & Biobehavioral Reviews, 144, 104978. 10.1016/j.neubiorev.2022.104978

German-Ponciano, L. J., Dutra Costa, B. P., Feitosa, L. M., Campos, K. dos S., da Silva Chaves, S. N., Cueto-Escobedo, J., Lima-Maximino, M., Rodríguez-Landa, J. F., & Maximino, C. (2020). Chrysin, but not flavone backbone, decreases anxiety-like behavior in animal screens. Neurochemistry International, 140, 104850. 10.1016/j.neuint.2020.104850

Germán-Ponciano, L. J., Puga-Olguín, A., Rovirosa-Hernández, M. D. J., Caba, M., Meza, E., & Rodríguez-Landa, J. F. (2020). Differential effects of acute and chronic treatment with the flavonoid chrysin on anxiety-like behavior and Fos immunoreactivity in the lateral septal nucleus in rats. Acta Pharmaceutica, 70(3), 387–397. 10.2478/acph-2020-0022

German-Ponciano, L. J., Rosas-Sánchez, G. U., Rivadeneyra-Domínguez, E., & Rodríguez-Landa, J. F. (2018). Advances in the Preclinical Study of Some Flavonoids as Potential Antidepressant Agents. Scientifica, 2018, e2963565. 10.1155/2018/2963565

Gomez, A. F., Barthel, A. L., & Hofmann, S. G. (2018). Comparing the efficacy of benzodiazepines and serotonergic anti-depressants for adults with generalized anxiety disorder: A metaanalytic review. Expert Opinion on Pharmacotherapy, 19, 883–894.

Grundmann, O., Nakajima, J.-I., Kamata, K., Seo, S., & Butterweck, V. (2009). Kaempferol from the leaves of Apocynum venetum possesses anxiolytic activities in the elevated plus maze test in mice. Phytomedicine, 16, 295–302. 10.1016/j.phymed.2008.12.020

Hanrahan, J. R., Chebib, M., & Johnston, G. A. R. (2011). Flavonoid modulation of GABA _A_ receptors. British Journal of Pharmacology, 163, 234–245. 10.1111/j.1476-5381.2011.01228.x

Hartley, D. E., Edwards, J. E., Spiller, C. E., Alom, N., Tucci, S., Seth, P., Forsling, M. L., & File, S. E. (2003). The soya isoflavone content of rat diet can increase anxiety and stress hormone release in the male rat. Psychopharmacology, 167(1), 46–53. 10.1007/s00213-002-1369-7

Hernandez-Leon, A., González-Trujano, M. E., & Fernández-Guasti, A. (2017). The anxiolytic-like effect of rutin in rats involves GABAA receptors in the basolateral amygdala. Behavioural Pharmacology, 28, 303–312. 10.1097/FBP.0000000000000290

Hooijmans, C. R., Tillema, A., Leenaars, M., & Ritskes-Hoitinga, M. (2010). Enhancing search efficiency by means of a search filter for finding all studies on animal experimentation in PubMed. Laboratory Animals, 44, 170–175. 10.1258/la.2010.009117

Hritcu, L., Ionita, R., Postu, P. A., Gupta, G. K., Turkez, H., Lima, T. C., Carvalho, C. U. S., & de Sousa, D. P. (2017). Antidepressant Flavonoids and Their Relationship with Oxidative Stress. Oxidative Medicine and Cellular Longevity, 2017, e5762172. 10.1155/2017/5762172

Huen, M. S. Y., Hui, K.-M., Leung, J. W. C., Sigel, E., Baur, R., Wong, J. T.-F., & Xue, H. (2003). Naturally occurring 2’-hydroxyl-substituted flavonoids as high-affinity benzodiazepine site ligands. Biochemical Pharmacology, 66, 2397–2407. 10.1016/j.bcp.2003.08.016

Hui, K. M., Huen, M. S. Y., Wang, H. Y., Zheng, H., Sigel, E., Baur, R., Ren, H., Li, Z. W., Wong, J. T.-F., & Xue, H. (2002). Anxiolytic effect of wogonin, a benzodiazepine receptor ligand isolated from *Scutellaria baicalensis* Georgi. Biochemical Pharmacology, 64, 1415–1424. 10.1016/s0006-2952(02)01347-3

Jäger, A. K., & Saaby, L. (2011). Flavonoids and the CNS. Molecules, 16, Artigo 2. 10.3390/molecules16021471

Jamal, H., Ansari, W. H., & Rizvi, S. J. (2008). Evaluation of chalcones—A flavonoid subclass, for, their anxiolytic effects in rats using elevated plus maze and open field behaviour tests. Fundamental & Clinical Pharmacology, 22, 673–681. 10.1111/j.1472-8206.2008.00639.x

Jia, S., Hou, Y., Wang, D., & Zhao, X. (2023). Flavonoids for depression and anxiety: A systematic review and meta-analysis. Critical Reviews in Food Science and Nutrition, 9, 8839–8849.

Kahnberg, P., Lager, E., Rosenberg, C., Schougaard, J., Camet, L., Sterner, O., Nielsen, E. Ø., Nielsen, M., & Liljefors, T. (2002). Refinement and Evaluation of a Pharmacophore Model for Flavone Derivatives Binding to the Benzodiazepine Site of the GABA _A_ Receptor. Journal of Medicinal Chemistry, 45, 4188–4201. 10.1021/jm020839k

Karim, N., Curmi, J., Gavande, N., Johnston, G. A., Hanrahan, J. R., Tierney, M. L., & Chebib, M. (2012). 2’-Methoxy-6-methylflavone: A novel anxiolytic and sedative with subtype selective activating and modulating actions at GABA(A) receptors. British Journal of Pharmacology, 165(4), 880–896. 10.1111/j.1476-5381.2011.01604.x

Karim, N., Gavande, N., Wellendorph, P., Johnston, G. A. R., Hanrahan, J. R., & Chebib, M. (2011). 3-Hydroxy-2’-methoxy-6-methylflavone: A potent anxiolytic with a unique selectivity profile at GABA(A) receptor subtypes. Biochemical Pharmacology, 82, 1971–1983. 10.1016/j.bcp.2011.09.002

Kumar, D., & Bhat, Z. A. (2014). Apigenin 7-glucoside from <IStachys tibetica Vatke and its anxiolytic effect in rats. Phytomedicine, 21, 1010–1014. 10.1016/j.phymed.2013.12.001

Kysil, E. V., Meshalkina, D. A., Frick, E. E., Echevarria, D. J., Rosemberg, D. B., Maximino, C., Lima, M. G., Abreu, M. S., Giacomini, A. C., Barcellos, L. J. G., Song, C., & Kalueff, A. V. (2017). Comparative Analyses of Zebrafish Anxiety-Like Behavior Using Conflict-Based Novelty Tests. Zebrafish, 14(3), 197–208. 10.1089/zeb.2016.1415

Li, J., Liu, Q.-T., Chen, Y., Liu, J., Shi, J.-L., Liu, Y., & Guo, J.-Y. (2016). Involvement of 5-HT1A Receptors in the Anxiolytic-Like Effects of Quercitrin and Evidence of the Involvement of the Monoaminergic System. Evidence-Based Complementary and Alternative Medicine: eCAM, 2016, 6530364. 10.1155/2016/6530364

Li, Y., Chen, Y., Chen, J., & Shen, C. (2023). Flavonoid metabolites in tea plant (*Camellia sinensis*) stress response: Insights from bibliometric analysis. Plant Physiology and Biochemistry, 202, 107934. 10.1016/j.plaphy.2023.107934

Liu, J., Zhai, W.-M., Yang, Y.-X., Shi, J.-L., Liu, Q.-T., Liu, G.-L., Fang, N., Li, J., & Guo, J.-Y. (2015). GABA and 5-HT systems are implicated in the anxiolytic-like effect of spinosin in mice. Pharmacology Biochemistry & Behavior, 128, 41–49. 10.1016/j.pbb.2014.11.003

Liu, L., Jia, L., Jian, P., Zhou, Y., Zhou, J., Wu, F., & Tang, Y. (2020). The Effects of Benzodiazepine Use and Abuse on Cognition in the Elders: A Systematic Review and Meta-Analysis of Comparative Studies. Frontiers in Psychiatry, 11. 10.3389/fpsyt.2020.00755

Liu, Z., Silva, J., Shao, A. S., Liang, J., Wallner, M., Shao, X. M., Li, M., & Olsen, R. W. (2021). Flavonoid compounds isolated from Tibetan herbs, binding to GABAA receptor with anxiolytic property. Journal of Ethnopharmacology, 267, 113630. 10.1016/j.jep.2020.113630

Longo, L. P., & Johnson, B. (2000). Addiction: Part I. Benzodiazepines—Side Effects, Abuse Risk and Alternatives. American Family Physician, 61, 2121–2128.

López-Rubalcava, C., & Estrada-Camarena, E. (2016). Mexican medicinal plants with anxiolytic or antidepressant activity: Focus on preclinical research. Journal of Ethnopharmacology, 186, 377–391. 10.1016/j.jep.2016.03.053

Marder, M., Estiú, G., Blanch, L. B., Viola, H., Wasowski, C., Medina, J. H., & Paladini, A. C. (2001). Molecular modeling and QSAR analysis of the interaction of flavone derivatives with the benzodiazepine binding site of the GABAA receptor complex. Biorganic & Medicinal Chemistry, 9, 323–335.

Marder, M., Viola, H., Wasowski, C., Fernández, S., Medina, J. H., & Paladini, A. C. (2003). 6-Methylapigenin and hesperidin: New valeriana flavonoids with activity on the CNS. *Pharmacology*, Biochemistry & Behavior, 75, 537–545. 10.1016/S0091-3057(03)00121-7

Martinez, M., & Sá, C. (2020). Highly Cited in the South: International Collaboration and Research Recognition Among Brazil’s Highly Cited Researchers. Journal of Studies in International Education, 24, 39–58. 10.1177/1028315319888890

Mendes, F. R. S., da Silva, A. W., Ferreira, M. K. A., Rebouças, E. de L., Moura Barbosa, I., da Rocha, M. N., Henrique Ferreira Ribeiro, W., Menezes, R. R. P. P. B. de, Magalhães, E. P., Marinho, E. M., Marinho, M. M., Bandeira, P. N., de Menezes, J. E. S. A., Marinho, E. S., & dos Santos, H. S. (2023). GABAA and serotonergic receptors participation in anxiolytic effect of chalcones in adult zebrafish. Journal of Biomolecular Structure and Dynamics, 41, 12426–12444. 10.1080/07391102.2023.2167116

Nachammai, V., Jeyabalan, S., & Muthusamy, S. (2021). Anxiolytic effects of silibinin and naringenin on zebrafish model: A preclinical study. Indian Journal of Pharmacology, 53, 457–464. 10.4103/ijp.IJP_18_20

Panche, A. N., Diwan, A. D., & Chandra, S. R. (2016). Flavonoids: An overview. Journal of Nutritional Science, 5, e47. 10.1017/jns.2016.41

Perez-Vizcaino, F., & Fraga, C. G. (2018). Research trends in flavonoids and health. Archives of Biochemistry and Biophysics, 646, 107–112. 10.1016/j.abb.2018.03.022

Pineda, P., Gregorutti, G., & Streitwieser, B. (2020). Emerging Decolonialized Research Collaboration: The Max Planck Society and the Leibniz Association in Latin America. Journal of Studies in International Education, 24, 59–78. 10.1177/1028315319888891

Qiu, Z.-K., Zhong, D.-S., He, J.-L., Liu, X., Chen, J.-S., & Nie, H. (2018). The anxiolytic-like effects of puerarin are associated with the changes of monoaminergic neurotransmitters and biosynthesis of allopregnanolone in the brain. Metabolic Brain Disease, 33(1), 167–175. 10.1007/s11011-017-0127-9

Richter, S. H., Garner, J. P., & Würbel, H. (2009). Environmental standardization: Cure or cause of poor reproducibility in animal experiments? Nature methods, 6(4), 257–261. 10.1038/nmeth.1312

Ritz, C., Baty, F., Streibig, J. C., & Gerhard, D. (2015). Dose-Response Analysis Using R. PLOS ONE, 10, e0146021. 10.1371/journal.pone.0146021

Rodrigues, E., Tabach, R., Galduróz, J. C. F., & Negri, G. (2008). Plants With Possible Anxiolytic and/or Hypnotic Effects Indicated by Three Brazilian Cultures—Indians, Afro-Brazilians, and River-Dwellers. Em Atta-ur-Rahman (Org.), Studies in Natural Products Chemistry (Vol. 35, p. 549–595). Elsevier. 10.1016/S1572-5995(08)80014-2

Rodríguez-Landa, J. F., German-Ponciano, L. J., Puga-Olguín, A., & Olmos-Vázquez, O. J. (2022). Pharmacological, Neurochemical, and Behavioral Mechanisms Underlying the Anxiolytic-and Antidepressant-like Effects of Flavonoid Chrysin. Molecules, 27, Artigo 11. 10.3390/molecules27113551

Rodríguez-Landa, J. F., Guillén-Ruiz, G., Hernández-López, F., Cueto-Escobedo, J., Rivadeneyra-Domínguez, E., Bernal-Morales, B., & Herrera-Huerta, E. V. (2021). Chrysin reduces anxiety-like behavior through actions on GABAA receptors during metestrus-diestrus in the rat. Behavioural Brain Research, 397, 112952. 10.1016/j.bbr.2020.112952

Rodríguez-Landa, J. F., Hernández-López, F., Martínez-Mota, L., Scuteri, D., Bernal-Morales, B., & Rivadeneyra-Domínguez, E. (2022). GABAA/Benzodiazepine Receptor Complex in the Dorsal Hippocampus Mediates the Effects of Chrysin on Anxiety-Like Behaviour in Female Rats. Frontiers in Behavioral Neuroscience, 15. 10.3389/fnbeh.2021.789557

Salgueiro, J. B., Ardenghi, P., Dias, M., Ferreira, M. B. C., Izquierdo, I., & Medina, J. H. (1997). Anxiolytic Natural and Synthetic Flavonoid Ligands of the Central Benzodiazepine Receptor Have No Effect on Memory Tasks in Rats. Pharmacology Biochemistry and Behavior, 58, 887–891. 10.1016/S0091-3057(97)00054-3

Samantha, A., Das, G., & Das, S. K. (2011). Roles of flavonoids in plants. International Journal of Pharmaceutical Sciences and Techonology, 6, 12–35.

Santos Oliveira, L., Kueirislene Amâncio Ferreira, M., Wagner de Queiroz Almeida-Neto, F., Wlisses da Silva, A., Ivo Lima Pinto Filho, J., Nunes da Rocha, M., Machado Marinho, E., Henrique Ferreira Ribeiro, W., Machado Marinho, M., Silva Marinho, E., Eire Silva Alencar de Menezes, J., & Dos Santos, H. S. (2024). Synthesis, molecular docking, ADMET, and evaluation of the anxiolytic effect in adult zebrafish of synthetic chalcone (E)-3-(4-(dimethylamino)phenyl)-1-(2-hydroxyphenyl)prop-2-en-1-one: An in vivo and in silico approach. Fundamental & Clinical Pharmacology, 38, 290–306. 10.1111/fcp.12960

Shanmugasundaram, J., Subramanian, V., Nadipelly, J., Kathirvelu, P., Sayeli, V., & Cheriyan, B. V. (2020). Anxiolytic-like activity of 5-methoxyflavone in mice with involvement of GABAergic and serotonergic systems—In vivo and in silico evidences. European Neuropsychopharmacology, 36, 100–110. 10.1016/j.euroneuro.2020.05.009

Shinfuku, M., Kishimoto, T., Uchida, H., Suzuki, T., Mimura, M., & Kikuchi, T. (2019). Effectiveness and safety of long-term benzodiazepine use in anxiety disorders: A systematic review and meta-analysis. International Clinical Psychopharmacology, 34, 211. 10.1097/YIC.0000000000000276

Skupien, S., & Rüffin, N. (2020). The Geography of Research Funding: Semantics and Beyond. Journal of Studies in International Education, 24(1), 24–38. 10.1177/1028315319889896

Spencer, J. P. E. (2007). The interactions of flavonoids within neuronal signalling pathways. Genes & Nutrition, 2, Artigo 3. 10.1007/s12263-007-0056-z

Stegenga, J. (2022). Evidence of effectiveness. Studies in History and Philosophy of Science, 91, 288–295. 10.1016/j.shpsa.2022.01.001

Tungmunnithum, D., Thongboonyou, A., Pholboon, A., & Yangsabai, A. (2018). Flavonoids and Other Phenolic Compounds from Medicinal Plants for Pharmaceutical and Medical Aspects: An Overview. Medicines, 5, Artigo 3. 10.3390/medicines5030093

Vesterinen, H. M., Sena, E. S., Egan, K. J., Hirst, T. C., Churolov, L., Currie, G. L., Antonic, A., Howells, D. W., & Macleod, M. R. (2014). Meta-analysis of data from animal studies: A practical guide. Journal of Neuroscience Methods, 221, 92–102. 10.1016/j.jneumeth.2013.09.010

Viechtbauer, W. (2005). Bias and Efficiency of Meta-Analytic Variance Estimators in the Random-Effects Model. Journal of Educational and Behavioral Statistics, 30, 261–293. 10.3102/10769986030003261

Viechtbauer, W. (2010). Conducting Meta-Analyses in R with the metafor Package. Journal of Statistical Software, 36, 1–48. 10.18637/jss.v036.i03

Vignes, M., Maurice, T., Lanté, F., Nedjar, M., Thethi, K., Guiramand, J., & Récasens, M. (2006). Anxiolytic properties of green tea polyphenol (-)-epigallocatechin gallate (EGCG). Brain Research, 1110, 102–115. 10.1016/j.brainres.2006.06.062

Viola, H., Wasowski, C., Levi de Stein, M., Wolfman, C., Silveira, R., Dajas, F., Medina, J. H., & Paladini, A. C. (1995). Apigenin, a component of *Matricaria recutita* flowers, is a central benzodiazepine receptors-ligand with anxiolytic effects. Planta Medica, 61, 213–216. 10.1055/s-2006-958058

Viola, H., Wolfman, C., Levi de Stein, M., Wasowski, C., Peña, C., Medina, J. H., & Paladini, A. C. (1994). Isolation of pharmacologically active benzodiazepine receptor ligands from *Tilia tomentosa* (Tiliaceae). Journal of Ethnopharmacology, 44, 47–53. 10.1016/0378-8741(94)90098-1

Vissiennon, C., Nieber, K., Kelber, O., & Butterweck, V. (2012). Route of administration determines the anxiolytic activity of the flavonols kaempferol, quercetin and myricetin— Are they prodrugs? The Journal of Nutritional Biochemistry, 23, 733–740. 10.1016/j.jnutbio.2011.03.017

Voelkl, B., Vogt, L., Sena, E. S., & Würbel, H. (2018). Reproducibility of preclinical animal research improves with heterogeneity of study samples. PLOS Biology, 16, e2003693. 10.1371/journal.pbio.2003693

Wang, C., Yang, S., Deng, J., Shi, L., Chang, J., Meng, J., Liu, W., Zeng, J., Xing, K., Wen, J., Liang, B., & Xing, D. (2023). The research progress on the anxiolytic effect of plant-derived flavonoids by regulating neurotransmitters. Drug Development Research, 84, 406–417. 10.1002/ddr.22038

Wang, H., Zhao, T., Liu, Z., Danzengquzhen Cisangzhuoma Ma, J., Li, X., Huang, X., & Li, B. (2023). The neuromodulatory effects of flavonoids and gut Microbiota through the gut-brain axis. Frontiers in Cellular and Infection Microbiology, 13, 1197646. 10.3389/fcimb.2023.1197646

Wasowski, C., & Marder, M. (2012). Flavonoids as GABAA receptor ligands: The whole story? Journal of Experimental Pharmacology, 4, 9. 10.2147/JEP.S23105

Wolfman, C., Viola, H., Paladini, A., Dajas, F., & Medina, J. H. (1994). Possible anxiolytic effects of chrysin, a central benzodiazepine receptor ligand isolated from Passiflora coerulea. Pharmacology, Biochemistry, and Behavior, 47, 1–4. 10.1016/0091-3057(94)90103-1

Xavier, J. da C., Almeida-Neto, F. W. Q., da Silva, P. T., Marinho, E. S., Ferreira, M. K. A., Magalhães, F. E. A., Nogueira, C. E. S., Bandeira, P. N., de Menezes, J. E. S. A., Teixeira, A. M. R., & Santos, H. S. dos. (2020). Structural characterization, electronic properties, and anxiolytic-like effect in adult zebrafish (*Danio rerio*) of cinnamaldehyde chalcone. Journal of Molecular Structure, 1222, 128954. 10.1016/j.molstruc.2020.128954

Yang, X., Fang, Y., Chen, H., Zhang, T., Yin, X., Man, J., Yang, L., & Lu, M. (2021). Global, regional and national burden of anxiety disorders from 1990 to 2019: Results from the Global Burden of Disease Study 2019. Epidemiology and Psychiatric Sciences, 30, e36. 10.1017/S2045796021000275

Youdim, K. A., Dobbie, M. S., Kuhnle, G., Proteggente, A. R., Abbott, N. J., & Rice-Evans, C. (2003). Interaction between flavonoids and the blood-brain barrier: In vitro studies. Journal of Neurochemistry, 85, 180–192. 10.1046/j.1471-4159.2003.01652.x

Zanoli, P., Avallone, R., & Baraldi, M. (2000). Behavioral characterisation of the flavonoids apigenin and chrysin. Fitoterapia, 71, S117–S123. 10.1016/s0367-326x(00)00186-6

Zarei, M., Sarihi, A., Ahmadimoghaddam, D., & Soltani, E. (2021). Effects of Intracerebroventricular Micro-injection of Kaempferol on Anxiety: Possible GABAergic Mechanism Involved. Avicenna Journal of Neuro Psycho Physiology, 8, 109–114. 10.32592/ajnpp.2021.8.2.108

