## Supplementary figures and images for "Flavonoids as anxiolytics in animal tests: Bibliometry and meta-analysis of the effects of flavonoids on anxiety-like behavior in animal tests"

### Figure S1

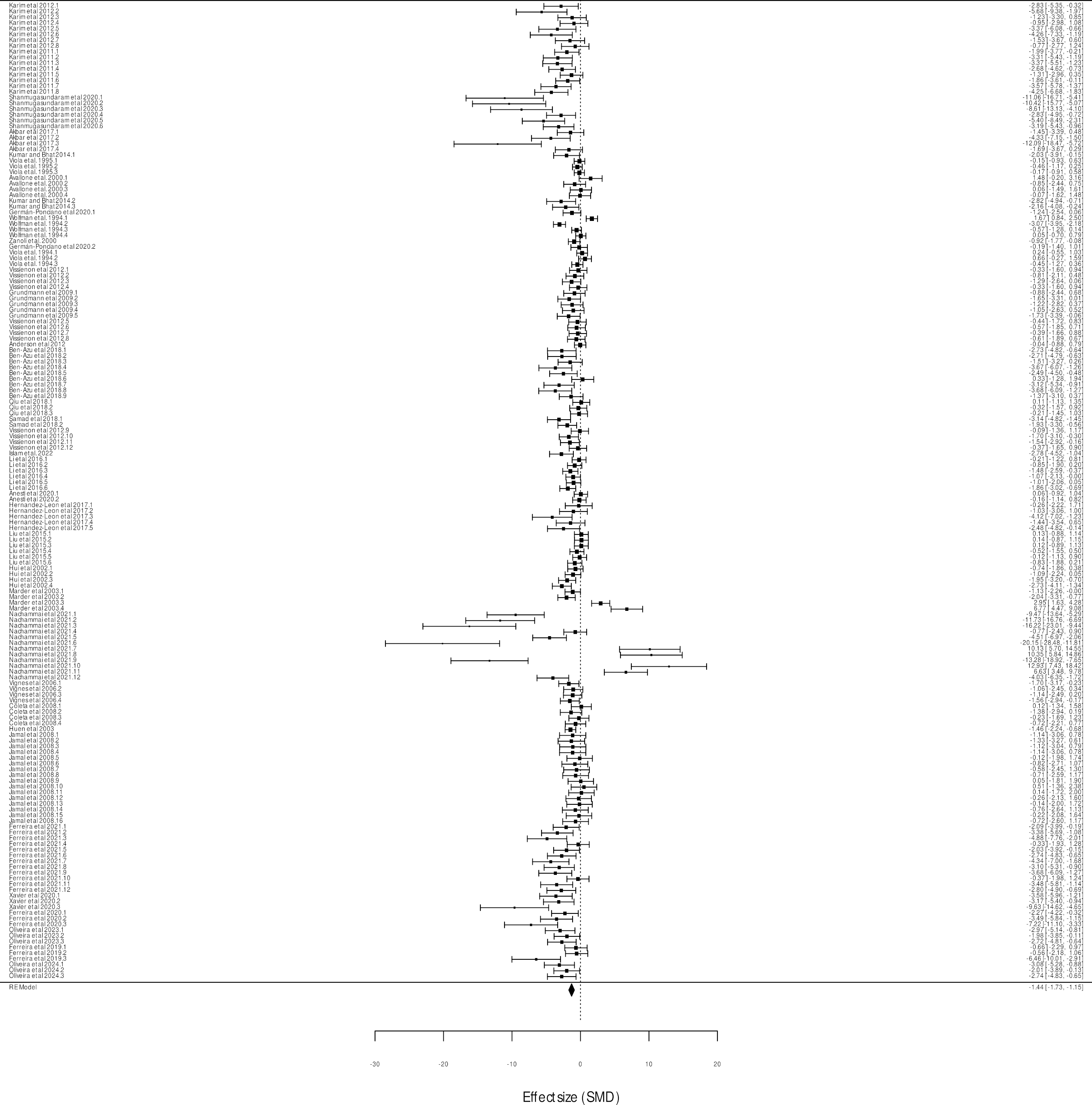

### Figure S2

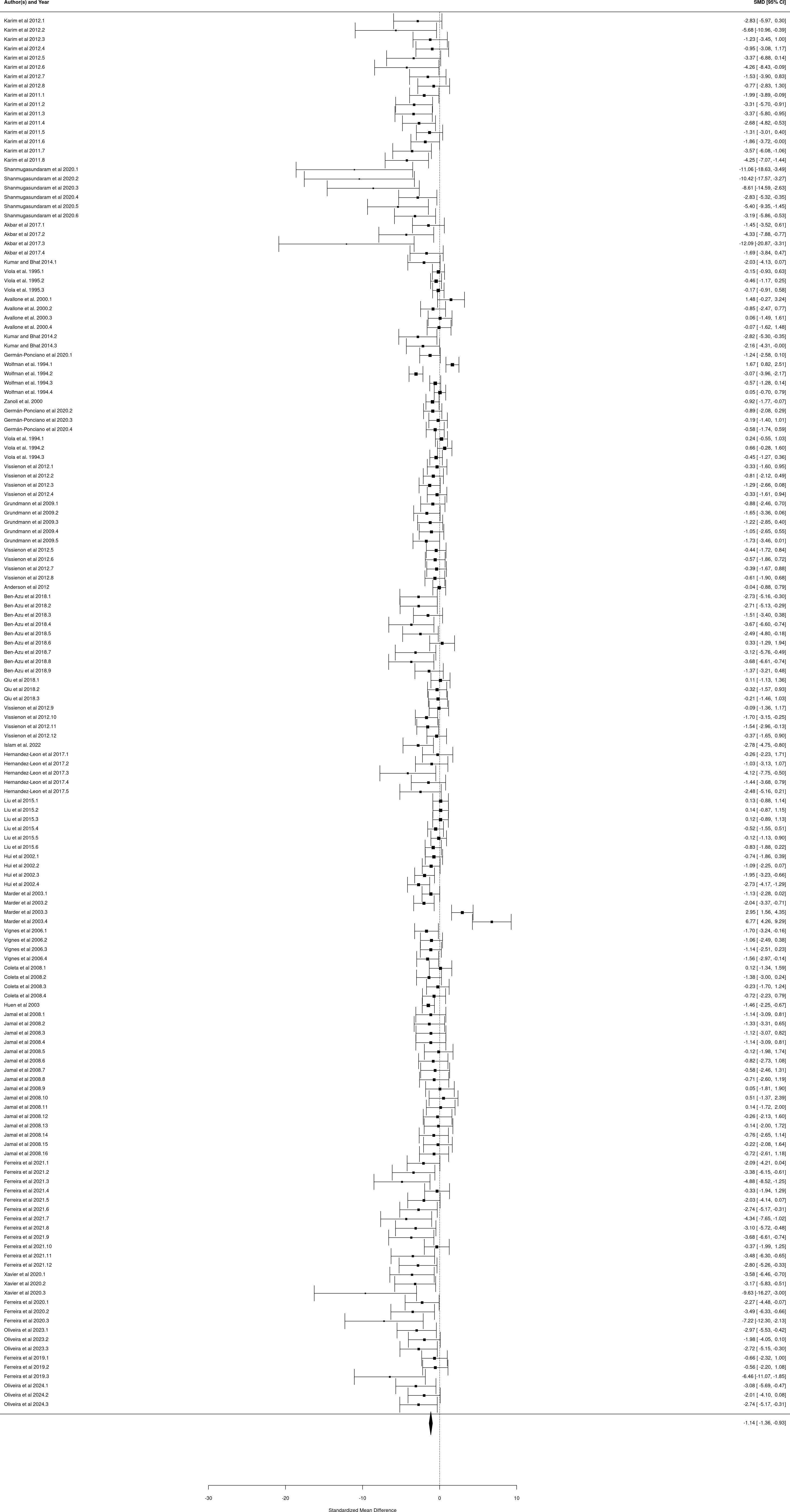

### Figure S3

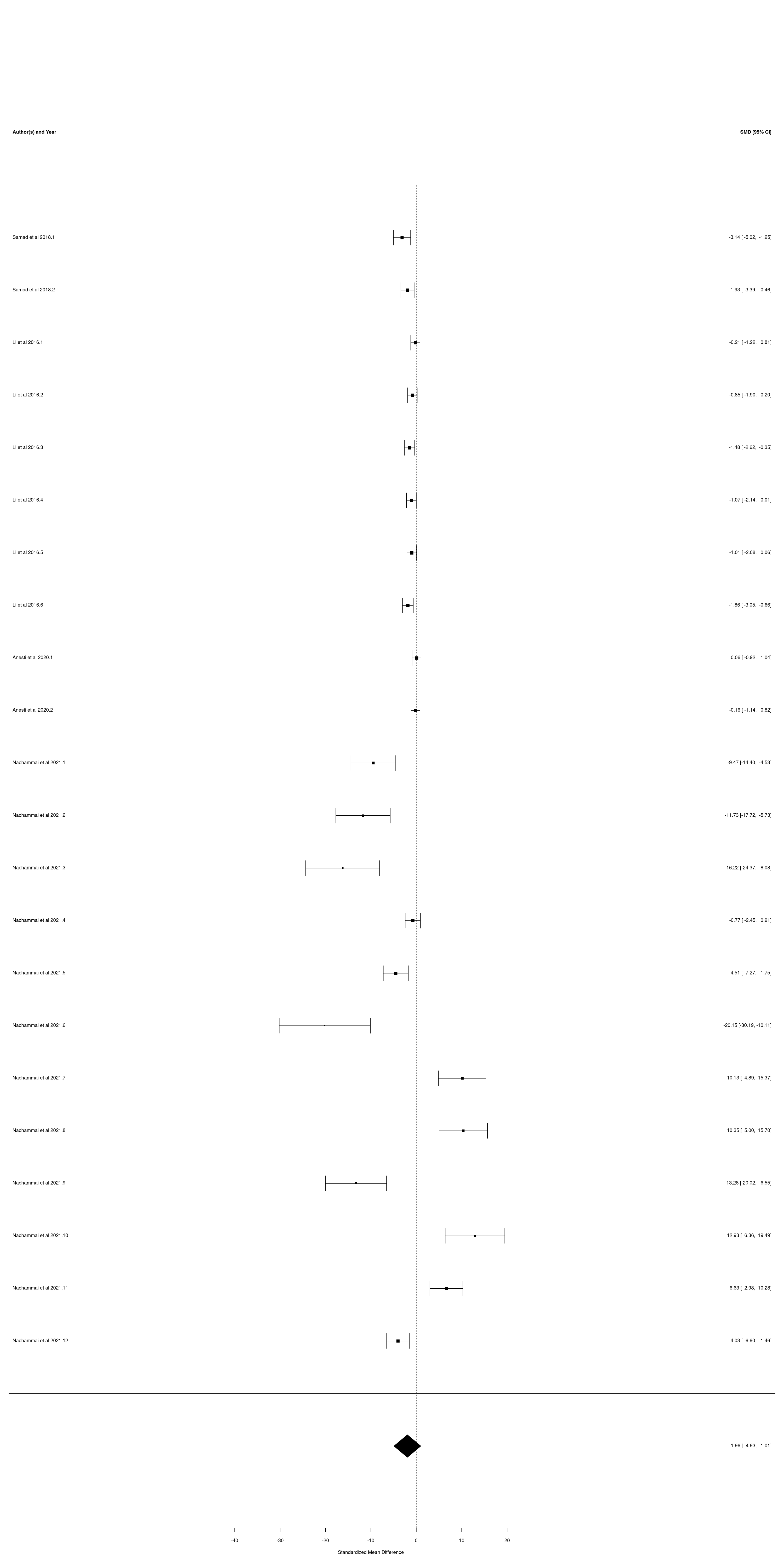
